# Deletion of two candidate depolymerases with overlapping genetic effects increases PHA yield in *Rhodopseudomonas palustris* TIE-1

**DOI:** 10.64898/2026.11.19.767437

**Authors:** Yantong Chen, Tahina Onina Ranaivoarisoa, Riyaz Ditter, Joshua AV Blodgett, Arpita Bose

## Abstract

Polyhydroxyalkanoate (PHA) is a microbial carbon-storage polymer with potential as a renewable bioplastic, but intracellular remobilization can limit polymer retention. The metabolically versatile bacterium *Rhodopseudomonas palustris* TIE-1 accumulates PHA during photoheterotrophic growth, yet the enzymes that control PHA turnover in this organism remain incompletely studied. We identified PhaZ2 (Rpal_0568) as a second candidate PHA depolymerase that is evolutionarily and architecturally distinct from the previously annotated PhaZ1 (Rpal_0578). Deleting either PhaZ1 or PhaZ2 alone did not substantially increase PHA content under tested growth conditions, whereas the double mutant accumulated 2.3-fold and 1.7-fold more PHA than wild type under non-nitrogen-fixing and nitrogen-fixing photoheterotrophic conditions, respectively, supporting overlapping roles in PHA turnover. All deletion strains showed longer doubling times but reached final culture densities similar to wild type. The double mutant showed its strongest retention phenotype under severe but growth-permissive nutrient limitation, including 3.7-fold more PHA than wild type at 33% phosphorus availability. A two-stage cultivation strategy that separated biomass formation from nutrient-limited storage raised volumetric PHA titer to 0.180 mg per mL culture in the double mutant, 1.5-fold more than wild type and approximately 3.4-fold higher than single-stage 33% phosphorus limitation and 4.0-fold higher than single-stage 8% nitrogen limitation (cross-experiment comparison), despite lower PHA content per unit biomass. Finally, an LC-MS-calibrated FTIR spectroscopy model predicted PHA content with an R² of 0.928 (*n* = 15) and reproduced the major biological trends. These results identify genetically redundant candidate depolymerases as targets for improving PHA retention in TIE-1 and establish a cultivation and screening framework for strain engineering.

**Importance:** Persistent pollution from petroleum-based plastics has intensified the need for biodegradable plastics produced from renewable resources. Polyhydroxyalkanoate (PHA) is a microbially synthesized bioplastic, but microbial cells can also consume the stored polymer, reducing its net yield. The metabolically versatile, photosynthetic purple non-sulfur bacterium (PNSB) *Rhodopseudomonas palustris* TIE-1 is a promising production organism because it can photosynthesize, assimilate atmospheric nitrogen, and utilize diverse carbon sources. We show that two candidate PHA depolymerases, PhaZ1 and PhaZ2, likely have overlapping roles in PHA turnover. Removing both candidates increased polymer retention, particularly when nitrogen or phosphorus was limited, and a two-stage cultivation strategy further increased PHA titer to 0.180 mg mL⁻¹ of culture. We also developed a rapid infrared spectroscopy method to track polymer content across strains and growth conditions. These findings offer complementary genetic, cultivation, and screening strategies for improving microbial bioplastic production and may help advance sustainable alternatives to petroleum-derived plastics.

## Introduction

Plastic pollution imposes major ecological and economic burdens, creating a need for sustainable alternatives to petroleum-based plastics. Microbially produced biodegradable plastics, such as polyhydroxyalkanoates (PHAs), have the potential to alleviate the environmental impact of conventional plastics (1,2). Nevertheless, the elevated cost of bioplastic production remains a major barrier to industrial-scale application, partially because many production organisms depend on organic carbon sources (3). *Rhodopseudomonas palustris* TIE-1 is a metabolically versatile, non-model purple non-sulfur bacterium (PNSB) that is capable of producing PHA. TIE-1 can utilize a broad range of carbon sources, including carbon dioxide and many organic substrates. It can also both assimilate ammonium salts or fix nitrogen from the atmosphere as the nitrogen source (4). This metabolic versatility renders it a useful organism for studying bioplastic production.

Although TIE-1 can synthesize PHA, stored PHA granules can also be remobilized, especially under starved conditions, reducing net PHA yield. PHA depolymerases, encoded by PhaZ, catalyze the degradation of stored PHA and are thus potential genetic targets for improving PHA accumulation (5). In *Rhodobacter sphaeroides*, deletion of an intracellular PhaZ depolymerase increased PHA accumulation under nitrogen-rich conditions (6). However, our previous work in TIE-1 showed that deletion of PhaZ1 (Rpal_0578) alone did not substantially enhance PHA production (5). This result suggested that PHA retention in TIE-1 may not be controlled by a single depolymerase gene, indicating that additional, functionally redundant depolymerase enzymes may contribute to PHA turnover (7). The TIE-1 genome contains a second *phaZ*-like locus, Rpal_0568, whose predicted product differs in domain architecture from PhaZ1. Functional redundancy between two such enzymes could explain why deleting *phaZ1* alone was insufficient to strongly increase PHA retention.

In addition to genetic factors, nutrient availability plays a critical role in regulating PHA accumulation and degradation. PHA storage is often stimulated when carbon is in excess relative to other growth-limiting nutrients (8). Specifically, nitrogen-to-carbon or phosphorus-to-carbon balance may influence whether carbon is allocated toward biomass accumulation or PHA storage, as PHA biosynthesis acts as the carbon storage sink (9,10). Moreover, the effect of PhaZ deletion may depend on growth stage. During active growth, PHA mobilization may be limited because carbon is being assimilated into biomass. In contrast, during stationary phase, PHA degradation may become more important for maintaining cellular metabolism, making depolymerase deletion more likely to enhance PHA retention (11–13).

Capturing these dynamic shifts across multiple growth phases and nutrient regimes requires efficient quantification; however, conventional PHA quantification methods often rely on chemical degradation of PHA polymers into crotonic acid, followed by analysis using GC or LC–MS (14,15). Although these methods provide quantitative measurements, they require extensive sample preparation, which limits the number of growth conditions and time points that can be analyzed. To address this limitation, we developed an FTIR-based approach for monitoring PHA accumulation in TIE-1. FTIR requires less sample processing and thus enables increased throughput that allows testing across strains, growth stages, and conditions. In addition, FTIR measures absorbance signals from intact dried biomass, including the PHA-associated carbonyl peak, rather than relying only on chemical degradation products. Therefore, an LC–MS-calibrated FTIR workflow can provide a faster screening method for estimating PHA accumulation while preserving LC–MS as the reference method for quantitative validation (15–17).

In this study, we examined the genetic basis of PHA retention in TIE-1. We identified PhaZ2 as a second candidate PHA depolymerase, constructed markerless *phaZ2* and *phaZ1 phaZ2* deletion mutants with homologous recombination (18–20), and compared growth and PHA accumulation across photoheterotrophic non nitrogen fixing and nitrogen fixing conditions (5,21). We further tested the effects of nutrient limitation, because restricting nitrogen or phosphorus relative to carbon sources is known to redirect metabolic flux toward PHA biosynthesis rather than biomass production (3). Under these conditions, cells experience an imbalance between carbon assimilation and biomass growth capacity, leading to increased accumulation of storage polymers such as PHA. In TIE-1, we tested PHA retention performance under nutrient limitation, and a two-stage cultivation strategy in which cells first accumulated biomass under nutrient-replete conditions and were then transferred to nutrient-limited and carbon-replete medium to promote PHA storage. We further show that FTIR spectroscopy calibrated by LC-MS provides a higher-throughput method for screening PHA accumulation across growth stages and cultivation conditions. Together, this work identifies functionally redundant depolymerase activity as a regulator of PHA retention in TIE-1 and provides genetic, cultivation, and analytical strategies for improving bioplastic production in purple non-sulfur bacteria.

## Results

### PhaZ1 and PhaZ2 are evolutionarily distinct PHA depolymerase candidates

PhaZ2 was identified by searching the TIE-1 genome for homologues of characterized and annotated PHA depolymerases. Rpal_0568 encodes a protein whose homologues are annotated as candidate PHA depolymerases in several other bacteria, and we designated it PhaZ2. To determine whether PhaZ1 and PhaZ2 represent evolutionarily distinct enzymes, we compared both with representative bacterial PHA depolymerases by multiple sequence alignment and phylogenetic analysis. The resulting neighbor-joining protein tree separated the two candidates into distinct clades: PhaZ1 grouped with putative intracellular short-chain-length PHA depolymerases, whereas PhaZ2 grouped with putative extracellular catalytic-domain type 1 short-chain-length PHA depolymerases (Fig. 1A). Accordingly, PhaZ1 contains a DepA/PHA depolymerase-like domain spanning most of the protein, consistent with other PHA depolymerases in the family (Fig. 1C), whereas PhaZ2 contains a CE1/LpqC esterase-like hydrolase domain.

**Figure 1.**
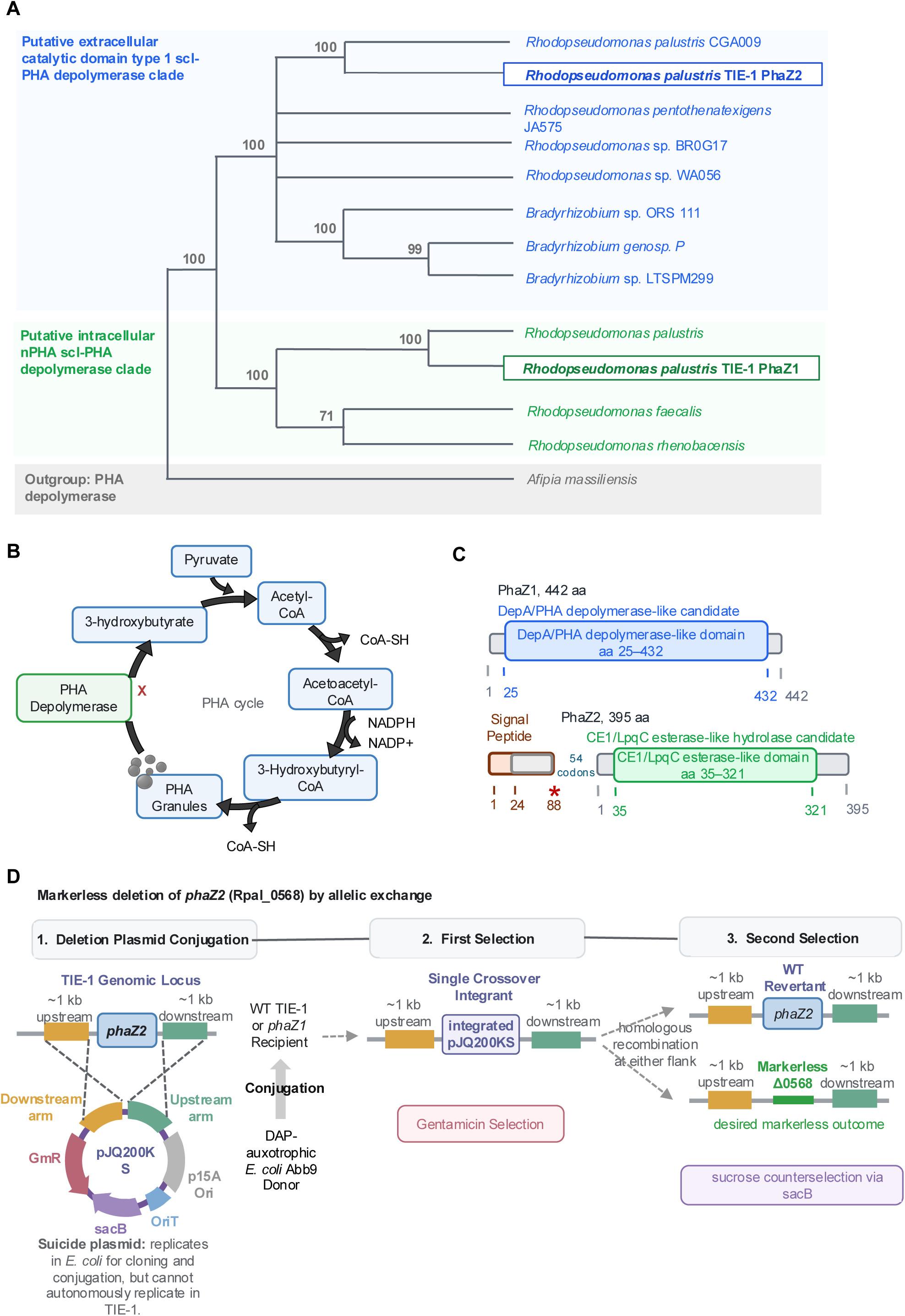
Predicted roles, evolutionary relationships, and deletion schematic of the TIE-1 PhaZ candidates. (A) Neighbor-joining tree of 13 representative bacterial PHA depolymerase sequences. Evolutionary distances were calculated using the p-distance method after partial deletion of positions with less than 95% site coverage, leaving 370 aligned positions. Branch support was assessed using 1,000 bootstrap replicates; bootstrap percentages are shown at supported nodes. PhaZ2 and PhaZ1 are highlighted within the putative extracellular and intracellular short-chain-length PHA depolymerase clades, respectively. *Afipia massiliensis* PHA depolymerase was used as the outgroup. Scale bar: 0.2 substitutions per site. (B) Proposed PHA synthesis and turnover pathway in *Rhodopseudomonas palustris* TIE-1, showing carbon flow from acetyl-CoA to PHA and the predicted block in PHA depolymerization after PhaZ deletion. (C) Predicted domain architectures of the TIE-1 PhaZ candidates. PhaZ1 (Rpal_0578; 442 amino acids) contains a DepA/PHA depolymerase-like domain spanning residues 25–432. PhaZ2 (Rpal_0568; 395 amino acids) contains a CE1/LpqC esterase-like hydrolase domain spanning residues 35–321. An 88-codon open reading frame lies immediately upstream of *phaZ2* and encodes a predicted Sec/SPI signal peptide at residues 1–24 (Fig. S3). This reading frame terminates at an in-frame stop codon (asterisk), after which 54 codons precede the *phaZ2* start codon. The signal peptide and the depolymerase coding sequence therefore cannot be translated as single polypeptide. The stop codon was confirmed by whole-genome sequencing of TIE-1 in this study. Domain boundaries were assigned by search against the NCBI Conserved Domain Database. (D) Approximately 1 kb upstream and downstream of Rpal_0568 were assembled into the suicide vector pJQ200KS and introduced into wild-type TIE-1 or the ΔRpal_0578 background by conjugation from a diaminopimelic-acid-auxotrophic Escherichia coli donor. Gentamicin selection isolated single-crossover integrants, and sacB-mediated sucrose counterselection resolved the integrated plasmid to yield either a wild-type revertant or the markerless deletion. The same workflow generated ΔRpal_0568 and ΔRpal_0578 ΔRpal_0568 strains.

The region immediately upstream of the annotated Rpal_0568 start encodes a canonical Sec/SPI signal peptide, which SignalP 6.0 assigned to the Sec/SPI class with a likelihood of 0.999, with predicted cleavage between positions 24 and 25 (Fig. S3). However, this signal peptide is separated from the annotated coding sequence by an in-frame stop codon, which we confirmed in our own whole-genome sequencing of TIE-1 (BioProject PRJNA1541609) rather than inferring from the reference sequence. The signal peptide and the Rpal_0568 coding sequence therefore cannot be translated as a single polypeptide. SignalP analysis of the Rpal_0568 product alone predicts no secretion signal (OTHER class, likelihood 1.00; Fig. S3), so PhaZ2 likely remains intracellular despite grouping with homologues annotated as extracellular short-chain-length PHA depolymerases.

### Marker-less gene deletion mutants were constructed to test candidate depolymerase function

To test whether PhaZ2 contributes to PHA metabolism independently of or in concert with PhaZ1, we generated a panel of test strains consisting of wild-type TIE-1, Δ*phaZ1*, Δ*phaZ2*, and the double mutant Δ*phaZ1* Δ*phaZ2* (Table 1). This panel allowed the phenotypes of the single mutants and the double mutant to be compared against the proposed PHA synthesis and turnover pathway shown in Fig. 1B.

**Table 1.**
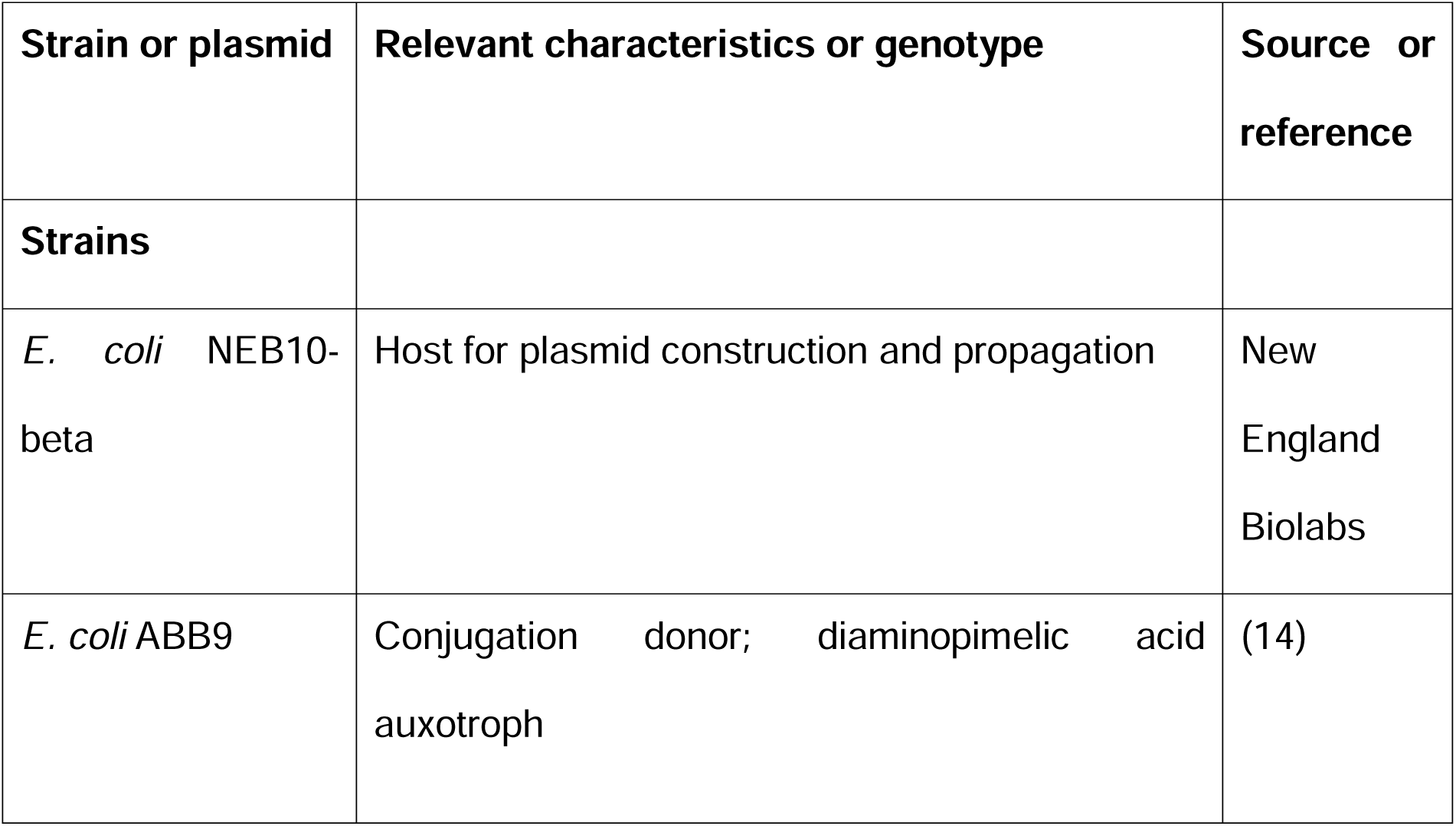
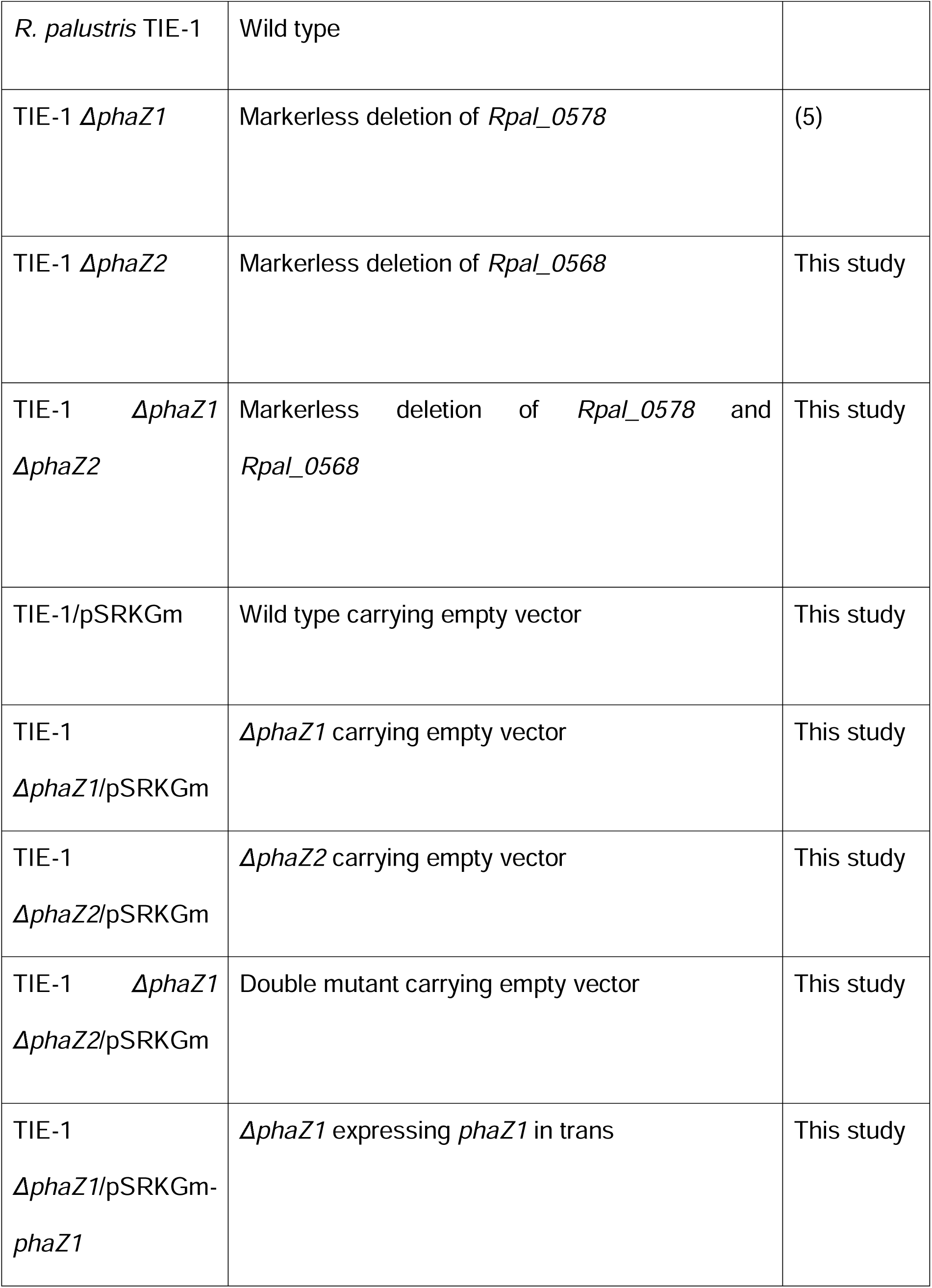
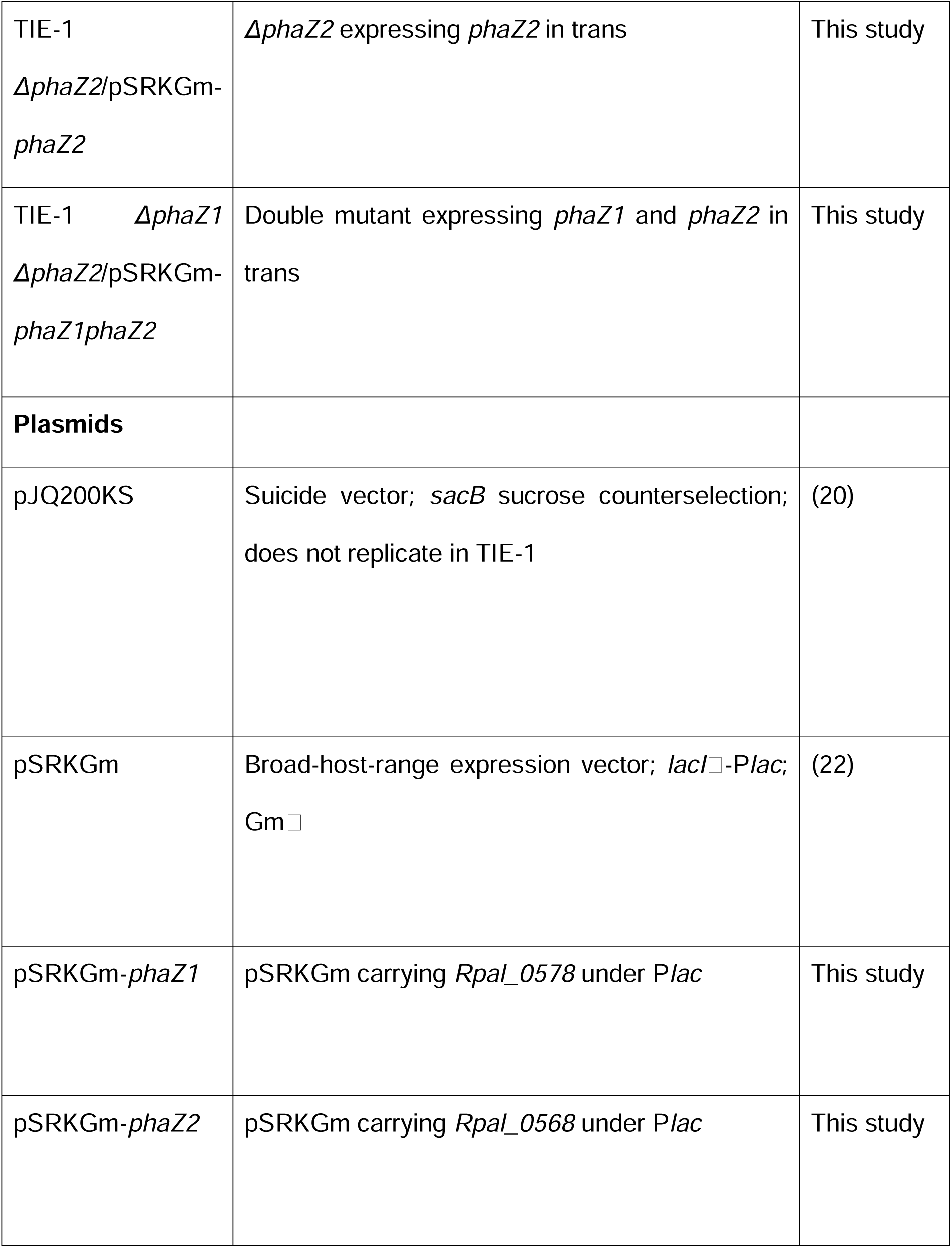
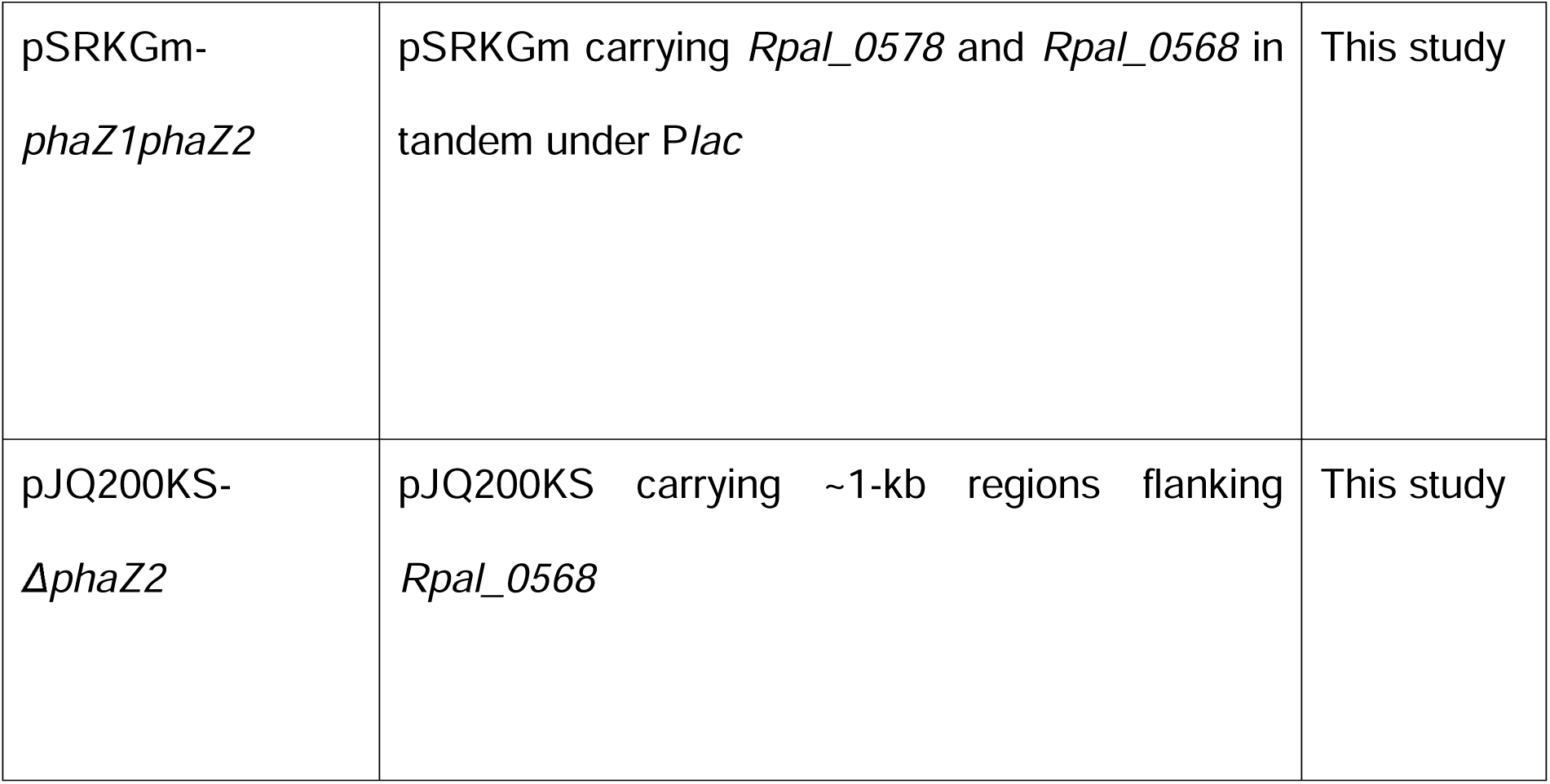
Strains and Plasmids used in this study.

| Strain or plasmid | Relevant characteristics or genotype | Source or reference |
|---|---|---|
| **Strains** |  |  |
| *E. coli* NEB10-beta | Host for plasmid construction and propagation | New England Biolabs |
| *E. coli* ABB9 | Conjugation donor; diaminopimelic acid auxotroph | (14) |
| *R. palustris* TIE-1 | Wild type |  |
| TIE-1 *ΔphaZ1* | Markerless deletion of *Rpal_0578* | (5) |
| TIE-1 *ΔphaZ2* | Markerless deletion of *Rpal_0568* | This study |
| TIE-1 *ΔphaZ1* *ΔphaZ2* | Markerless deletion of *Rpal_0578* and *Rpal_0568* | This study |
| TIE-1/pSRKGm | Wild type carrying empty vector | This study |
| TIE-1 *ΔphaZ1*/pSRKGm | *ΔphaZ1* carrying empty vector | This study |
| TIE-1 *ΔphaZ2*/pSRKGm | *ΔphaZ2* carrying empty vector | This study |
| TIE-1 *ΔphaZ1* *ΔphaZ2*/pSRKGm | Double mutant carrying empty vector | This study |
| TIE-1 *ΔphaZ1*/pSRKGm-*phaZ1* | *ΔphaZ1* expressing *phaZ1* in trans | This study |
| TIE-1 *ΔphaZ2*/pSRKGm-*phaZ2* | *ΔphaZ2* expressing *phaZ2* in trans | This study |
| TIE-1 *ΔphaZ1* *ΔphaZ2*/pSRKGm-*phaZ1phaZ2* | Double mutant expressing *phaZ1* and *phaZ2* in trans | This study |
| **Plasmids** |  |  |
| pJQ200KS | Suicide vector; *sacB* sucrose counterselection; does not replicate in TIE-1 | (20) |
| pSRKGm | Broad-host-range expression vector; *lacI*[illegible]-P*lac*; Gm[illegible] | (22) |
| pSRKGm-*phaZ1* | pSRKGm carrying *Rpal_0578* under P*lac* | This study |
| pSRKGm-*phaZ2* | pSRKGm carrying *Rpal_0568* under P*lac* | This study |
| pSRKGm-*phaZ1phaZ2* | pSRKGm carrying *Rpal_0578* and *Rpal_0568* in tandem under P*lac* | This study |
| pJQ200KS-*ΔphaZ2* | pJQ200KS carrying ~1-kb regions flanking *Rpal_0568* | This study |

To construct the Δ*phaZ*2 strain, we used a markerless allelic exchange strategy based on the suicide vector pJQ200KS (Fig. 1D). Approximately 1,000 bp upstream and downstream of phaZ2 were cloned into pJQ200KS to generate a deletion vector. The resulting plasmid was amplified and introduced into WT TIE-1, and single-crossover recombinants were isolated by gentamicin selection. Next, strains that underwent double-crossover events were selected with SacB-mediated sucrose counterselection. The same deletion strategy was applied in the Δ*phaZ*1 background to generate the Δ*phaZ1* Δ*phaZ2* double mutant.

Candidate mutants were screened by genomic PCR using primers flanking the PhaZ2 locus. The deletion strains produced PCR products consistent with the deletion of the PhaZ2 region, whereas wild-type TIE-1 produced the amplicon containing the locus (Fig. S1). Genomic sequencing of the strains further confirmed the intended markerless deletion in both the Δ*phaZ*2 and Δ*phaZ1* Δ*phaZ2* backgrounds. Together, these strains provided a defined genetic panel to test whether PHA retention in TIE-1 is controlled by PhaZ1, by PhaZ2, or by partially redundant activity of both candidate depolymerases.

### Deletion of *phaZ*1, *phaZ*2, or both created minor growth defects under non-nitrogen fixing and nitrogen fixing conditions

We compared growth and PHA accumulation in the depolymerase mutants under anoxic, photoheterotrophic conditions with butyrate as the sole carbon source (Fig. 2), since TIE-1 was shown to exhibit high PHA production under this condition (14,18). Growth was monitored by OD in Fresh Water medium containing 1 mM butyrate to exhibit clear growth patterns. Growth curves were fitted using a modified Gompertz model to quantitatively compare fitted doubling time and lag time (23). We also examined how nitrogen fixation affected growth by comparing non–nitrogen-fixing conditions, in which ammonium chloride (NH₄Cl) was supplied as a fixed nitrogen source, with nitrogen-fixing conditions, in which nitrogen gas served as the sole nitrogen source.

**Figure 2.**
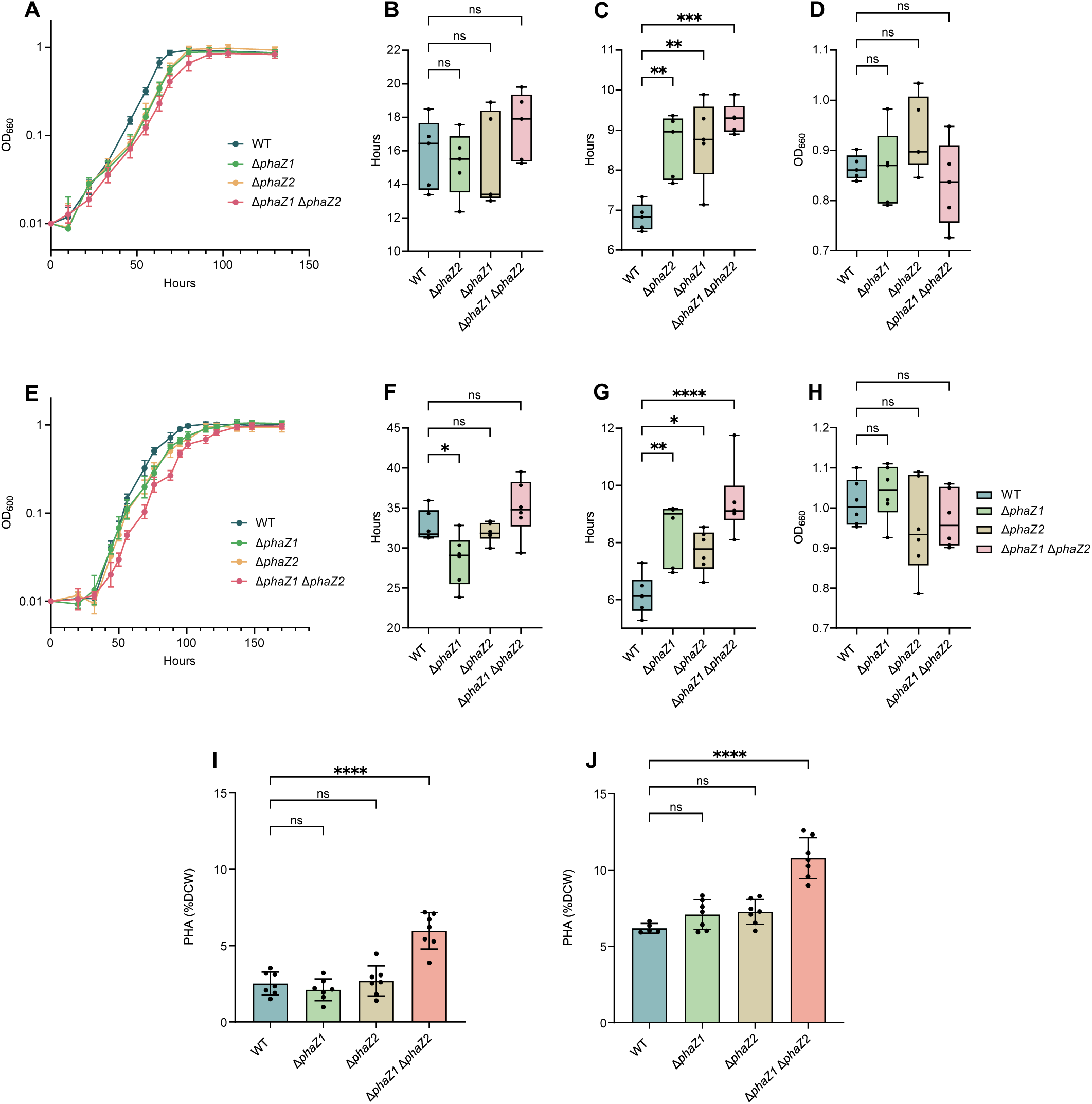
Growth and PHA accumulation of TIE-1 *phaZ* deletion strains under non-nitrogen-fixing and nitrogen-fixing photoheterotrophic conditions. (A - D) Growth, y axis is logarithmic, under non-nitrogen-fixing conditions with NH₄Cl supplied as fixed nitrogen: OD₆₆₀ time course (A), model-fitted lag time (B), doubling time (C), and final OD₆₆₀ (D). (E–H) Corresponding growth measurements under nitrogen-fixing conditions, in which N₂ was the sole nitrogen source: OD₆₆₀ time course (E), lag time (F), doubling time (G), and final OD₆₆₀ (H). Growth experiments used Freshwater medium containing 1 mM butyrate. (I, J) PHA content of wild type, *ΔphaZ1*, *ΔphaZ2*, and *ΔphaZ1 ΔphaZ2* under non-nitrogen-fixing (I) and nitrogen-fixing (J) conditions in freshwater medium containing 10 mM butyrate, harvested at early stationary phase. PHA was quantified by LC–MS and normalized to estimated dry cell weight. Growth parameters and PHA content were compared separately within each condition by ordinary one-way ANOVA followed by Tukey-adjusted comparisons of each deletion strain with wild type. Points represent biological replicates (*n* = 7 for PHA measurements, except *n* = 5 of wild type strains under nitrogen-fixing conditions; *n* = 5 for non-nitrogen-fixing growth and *n* = 6 for nitrogen-fixing growth); bars show the mean with standard deviation. ns, not significant; \**P <* 0.05; \*\**P <* 0.01; \*\*\**P <* 0.001; \*\*\*\**P <* 0.0001. The y-axis label for PHA quantification panels (I, J) is standardized as “PHA (% DCW)”; any panel-level label reading “PHB (DCW%)” should be interpreted as PHA (% DCW).

Under non-nitrogen-fixing conditions, no statistically significant difference in lag time or final OD were observed across the four variants. However, all three deletion mutant strains exhibited statistically longer model-fitted doubling time compared to WT, indicating that loss of the candidate depolymerases impose a traceable growth defect in the tested condition (*P =* 0.0014, 0.0026, 0.0001 respectively) (Fig. 2A-D). We observed a similar trend under nitrogen-fixing conditions. Growth curves and fitted growth parameters showed that the double mutant maintained measurable growth under nitrogen-fixing conditions, as the final OD and lag time show no statistical difference. However, all three knockout mutants showed statistically longer doubling times (*P =* 0.0018, 0.0309, <0.0001) (Fig. 2E-H).

### Single mutants did not significantly increase PHA retention photoheterotrophically, while the double mutant increased

PHA accumulation was measured under non-nitrogen-fixing photoheterotrophic conditions with 10 mM butyrate to accumulate sufficient biomass. Samples were collected at the early stationary phase around OD 2.0 and analyzed via LC-MS after acid digestion. Single deletion of either *phaZ1* or *phaZ2* did not strongly increase PHA (dry cell weight % wt/wt, DCW) retention relative to wild type, whereas the double mutant retained more PHA, ∼2.3x compared to WT (*P <* 0.0001) (Fig. 2I). We observed a similar trend under nitrogen-fixing photoheterotrophic condition with 10 mM butyrate. Both single mutants did not retain significantly higher PHA, but the double deletion mutant increased the PHA accumulation measurably, ∼1.7x compared to WT (*P <* 0.0001) (Fig. 2J). The pattern indicates that loss of one candidate depolymerase was insufficient to substantially increase PHA retention, while simultaneous removal of both candidates enhanced polymer retention.

Since PHA was measured as crotonic acid, we confirmed that the wild-type samples were above the limit of quantification. Crotonic acid concentrations ranged from 256 to 616 ppm for wild type and 678 to 1,270 ppm for the double mutants under non-nitrogen-fixing conditions, and from 779 to 876 ppm and 1,188 to 1,670 ppm respectively under nitrogen-fixing conditions, against a limit of quantification of 188 ppm for this curve. The lowest wild-type sample exceeded that limit by a factor of around 1.4. Only one Δ*phaZ2* sample at 161 ppm fell between the limits of detection and quantification, and neither its retention nor its exclusion alters any genotype comparison.

### Complementation did not restore growth defects in *phaZ* deletion strains

To test whether the growth and PHA retention phenotype of the Δ*phaZ1* Δ*phaZ2* double mutant is a result of loss of the deleted genes, we constructed a panel of seven strains carrying the broad-host-range inducible expression vector pSRKGm. Wild type, Δ*phaZ*1, Δ*phaZ*2, and Δ*phaZ1* Δ*phaZ2* strain received an empty vector, and each deletion strain additionally received a construct expressing the corresponding gene or genes in trans under a *lac* promoter (Table 1). All strains (*n*=4) were cultured in parallel under anoxic photoheterotrophic conditions at 25% nitrogen availability with an argon headspace, gentamicin, and IPTG (10 mM).

A modified Gompertz model was fitted to individual growth curves (*n* = 4), yielding lag time, maximum specific growth rate, and final optical density for each culture. We found that each parameter responded differently to complementation (Fig. 3A, C–E).

**Figure 3.**
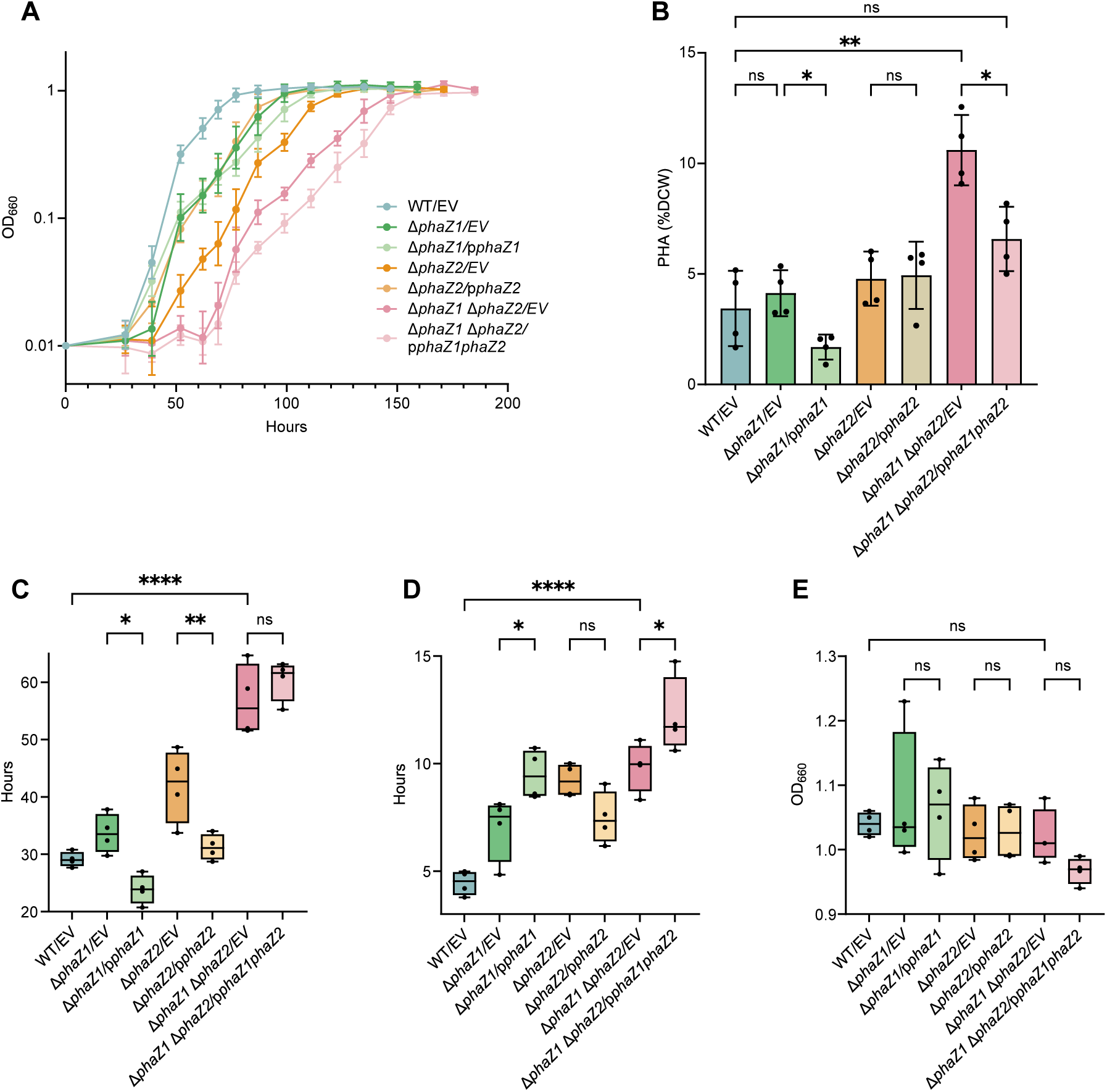
Genetic complementation of *phaZ1* and *phaZ2* partially restores PHA content and lag time but not growth rate. Wild type and each deletion strain carrying empty pSRKGm were compared with each deletion strain carrying pSRKGm expressing the corresponding gene or genes in trans under the vector IPTG-inducible *lac* promoter. All strains were grown anoxically under photoheterotrophic conditions in freshwater medium at 25% nitrogen availability with argon headspace, in the presence of gentamicin and IPTG. Strain labels are abbreviated: EV, empty pSRKGm vector; p*phaZ1*, pSRKGm-*phaZ1*; p*phaZ2*, pSRKGm-*phaZ2*; p*phaZ1phaZ2*, pSRKGm-*phaZ1phaZ2*. Full genotypes are given in Table 1.(A) OD₆₆₀ growth curve; the y axis is logarithmic. (B) PHA content as a percentage of dry cell weight, determined by LC–MS. (C–E) Growth parameters obtained by fitting a modified Gompertz model to each biological replicate individually: lag time (C), doubling time (D), and final OD₆₆₀ (E). PHA content was compared by Brown-Forsythe and Welch ANOVA with Dunnett T3 multiple comparisons, selected because group variances were unequal; growth parameters were compared by ordinary one-way ANOVA with Šídák-adjusted comparisons. Comparisons were specified in advance between each complemented strain and its corresponding empty-vector parent, and between *ΔphaZ1 ΔphaZ2* carrying empty vector and wild type. Points represent biological replicates (*n* = 4); bars show the mean with standard deviation. ns, not significant; *P < 0.05; **P < 0.01; ****P < 0.0001. Colors in panel A were selected from a colorblind-safe palette (Okabe-Ito; Wong, 2011) and each strain is additionally distinguished by a unique point shape.

Lag time in the double mutant was approximately two-fold longer than in wild type carrying empty vector (*P* < 0.0001). Expression of *phaZ1* in the ΔphaZ1 mutant shortened the lag phase by approximately 29% relative to the empty-vector parent (*P* = 0.0114). Likewise, expression of *phaZ2* in the ΔphaZ2 mutant shortened it by approximately 26% (*P* = 0.0054). In both single mutants, lag was restored to at or below the wild-type value. Expression of both genes in the double mutant did not further shorten lag (*P* = 0.6412).

Doubling time showed the opposite pattern for *phaZ1*-containing constructs. The double mutant grew approximately 2.2-fold more slowly than wild type (*P* < 0.0001). Expression of *phaZ1* in the Δ*phaZ1* mutant increased doubling time a further 1.4-fold relative to the empty-vector parent (*P* = 0.0344), whereas expression of *phaZ2* in the Δ*phaZ2* mutant had no significant effect (*P* = 0.2073). Expression of both genes in the double mutant also increased doubling time (*P* = 0.0495). Final optical density did not differ among any of the seven strains (all *P* ≥ 0.61).

### Complementation partially restored PHA retention in *phaZ* deletion strains

Wild type carrying empty vector accumulated 3.4 ± 1.7% DCW, compared with 10.6 ± 1.6% DCW in Δ*phaZ1* Δ*phaZ2* carrying empty vector (*P =* 0.0043), suggesting that the PHA retention phenotype in double mutant was reproduced under these conditions. Expression of both *phaZ1* and *phaZ2* in trans reduced PHA content in the double-mutant background to 6.6 ± 1.5% DCW (*P =* 0.0464), recovering ∼56% of the difference between the double mutant and wild type. PHA content in the complemented double mutant did not differ significantly from wild type carrying empty vector *(P = 0.1383)*. However, the confidence interval for this comparison (−0.96 to 7.25% DCW) is consistent with values ranging from full restoration to substantially incomplete restoration, so complete restoration cannot be inferred at this sample size (Fig. 3B). In addition, expression of *phaZ1* in the Δ*phaZ1* strain reduced PHA content relative to the empty-vector parent (*P = 0.0413*). However, expression of *phaZ2* in the Δ*phaZ2* strain produced no detectable change *(P > 0.999)*.

### Nutrient limitation increased PHA accumulation in TIE-1 *ΔphaZ1 ΔphaZ2*

We next tested whether nitrogen or phosphorus limitation would amplify the difference between wild type and Δ*phaZ1* Δ*phaZ2*, since limiting a nutrient other than carbon redirects surplus carbon toward PHA synthesis (8–10) and should therefore increase the pool available for remobilization. Since previous results suggest that only the double deletion strain increases PHA retention significantly, we grew WT and Δ*phaZ1* Δ*phaZ2* under different nitrogen and phosphorus availability conditions photoheterotrophically under 5 mM butyrate to monitor both the growth and PHA accumulation pattern.

For nitrogen-limitation experiments, WT and Δ*phaZ1* Δ*phaZ2* were compared under 100%, 25%, 8%, and 0% added NH₄Cl in Freshwater medium (0.3 g L⁻¹, 5.6 mM), representing nutrient-replete, moderately limited, strongly limited, and nitrogen-free conditions, respectively (Fig. 4A). To prevent nitrogen fixation from providing an alternative nitrogen source, the culture headspace was exchanged with argon gas. Growth remained near background under the 0% nitrogen condition. Thus, this negative-control condition was excluded from PHA quantification (Fig. 4B and C).

**Figure 4.**
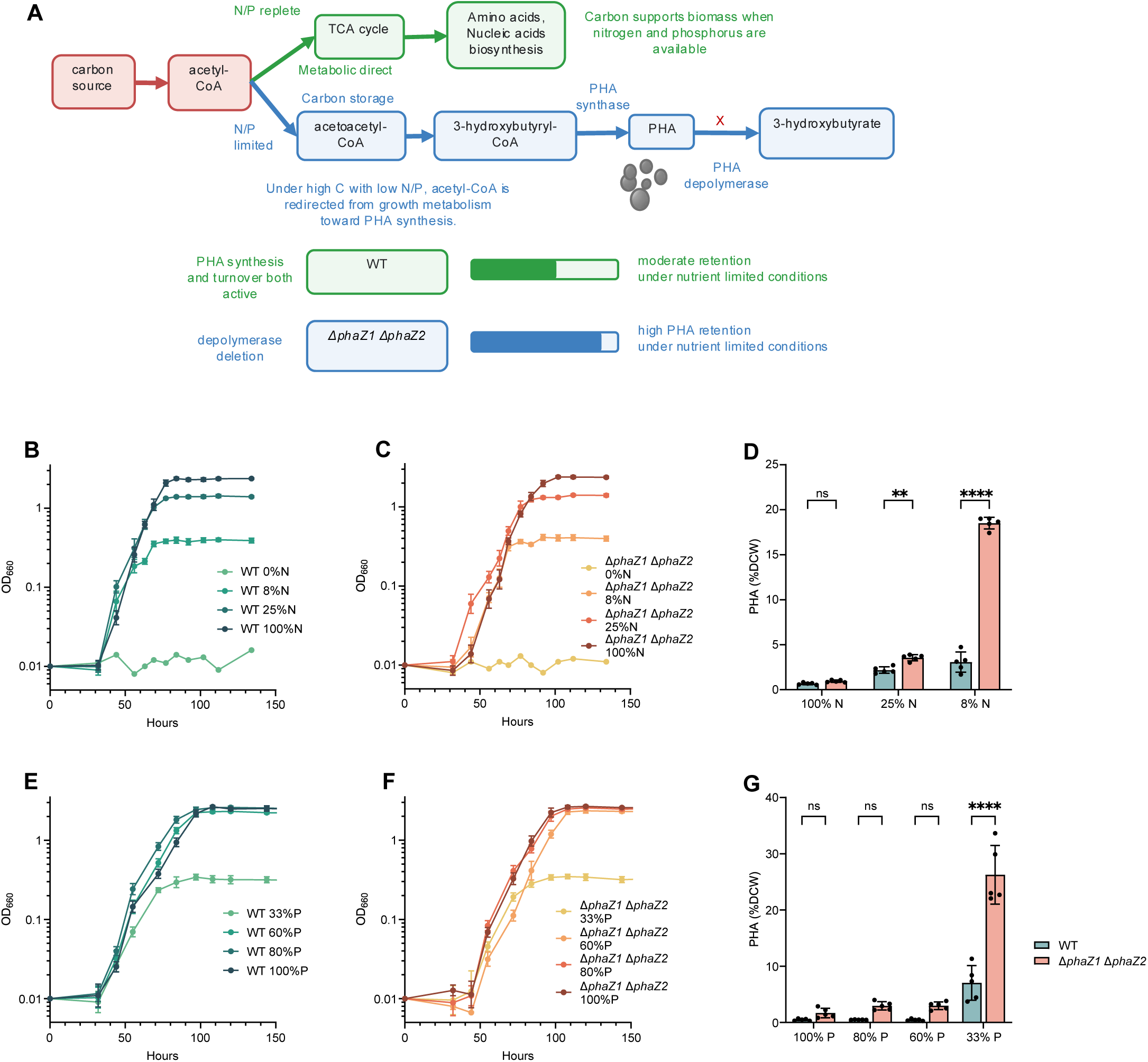
Nutrient availability modulates PHA retention in the *ΔphaZ1 ΔphaZ2* mutant. (A) Conceptual model of carbon allocation under nutrient-replete and nitrogen- or phosphorus-limited conditions. When nitrogen and phosphorus are available, carbon supports biosynthesis and growth; under limitation, excess carbon is redirected toward PHA storage. Loss of both candidate depolymerases is predicted to increase PHA retention. (B, C) Growth of WT (B) and *ΔphaZ1 ΔphaZ2* (C) under 100%, 25%, 8%, and 0% NH₄Cl availability, y axis is logarithmic. The headspace was exchanged with argon to prevent nitrogen fixation. (D) PHA content under the three nitrogen conditions that produced sufficient biomass for quantification; the 0% nitrogen condition was excluded. (E, F) Growth of WT (E) and *ΔphaZ1 ΔphaZ2* (F) under 100%, 80%, 60%, and 33% phosphorus availability. (G) PHA content under the corresponding phosphorus conditions. Cultures were grown with 5 mM butyrate and harvested 48 h after entering early stationary phase. PHA data were analyzed by ordinary two-way ANOVA with genotype and nutrient availability as fixed factors, including their interaction, followed by Šídák-adjusted comparisons between genotypes within each nutrient condition. Growth curves and bars show the mean with standard deviation; points represent biological replicates. ns, not significant; \*\**P <* 0.01; \*\*\*\**P <* 0.0001. Panels B–D correspond to nitrogen-limitation experiments; panels E–G correspond to phosphorus-limitation experiments. †FTIR-estimated PHA values in panels D and G that exceed the calibration maximum (3.89% DCW) are outside the validated calibration range; absolute values for these data points should be interpreted as estimates only.

Across the three quantifiable nitrogen conditions, PHA accumulation was significantly affected by nitrogen availability, genotype, and their interaction. Two-way ANOVA identified significant effects of nitrogen availability (*P <* 0.0001) and genotype (*P <* 0.0001), together with a significant genotype-by-nitrogen interaction (*P <* 0.0001). Nitrogen availability accounted for 47.65% of the total variation, whereas genotype and interaction accounted for 20.94% and 30.74%, respectively (Fig. 4D). The significant interaction indicated that the effect of *phaZ1 phaZ2* deletion depended strongly on nitrogen availability. PHA accumulation remained low under nitrogen-replete conditions and did not differ significantly between WT and Δ*phaZ1* Δ*phaZ2* after correction for multiple comparisons. In contrast, the double mutant accumulated significantly more PHA than WT under nitrogen-limited conditions. At 8% nitrogen availability, Δ*phaZ1* Δ*phaZ2* accumulated 18.51% PHA per dry cell weight (% DCW), compared with 3.06% DCW in WT, representing an approximately 6.1-fold increase. PHA content in the double mutant at 8% nitrogen was also approximately 28-fold higher than that of WT under nitrogen-replete conditions, which accumulated 0.67% DCW. Because the double-mutant PHA content at 8% nitrogen availability (18.51% DCW) falls well outside the FTIR calibration range (0.57–3.89% DCW), the LC-MS endpoint values reported in Data Set S1 should be considered the authoritative measurements for this high-accumulation condition.

A similar nutrient-dependent genotype effect was observed under phosphorus limitation. WT and Δ*phaZ1* Δ*phaZ2* grew under the 100%, 80%, 60%, and 33% phosphorus conditions, while growth declined with stronger phosphorus limitation. Growth under 5% phosphorus was insufficient to provide adequate biomass for PHA quantification; this condition was therefore excluded from the PHA analysis. Across the four quantifiable conditions, two-way ANOVA identified significant effects of phosphorus availability (*P <* 0.0001) and genotype (*P <* 0.0001), together with a significant genotype-by-phosphorus interaction (*P <* 0.0001). Phosphorus availability accounted for 60.75% of the total variation, whereas genotype and the interaction accounted for 14.23% and 19.59%, respectively. The significant interaction indicated that the effect of the double deletion varied with phosphorus availability (Fig. 4E-G).

At 100%, 80%, and 60% phosphorus availability, Δ*phaZ1* Δ*phaZ2 showed modest increases in PHA content* relative to WT, but these differences were not significant after correction for multiple comparisons. In contrast, under 33% phosphorus availability, the double mutant accumulated approximately 3.7-fold more PHA than WT under the same condition (adjusted *P <* 0.0001). Its PHA content under 33% phosphorus was also approximately 54.5-fold higher than that of WT under phosphorus-replete conditions. Together, these results show that the PHA-retention phenotype associated with combined *phaZ1 and phaZ2* deletion depended on nutrient availability and was strongest under severe but growth-permissive nitrogen or phosphorus limitation. Similarly, because the double-mutant PHA content at 33% phosphorus availability also substantially exceeds the FTIR calibration range, the LC-MS endpoint values in Data Set S1 provide the authoritative measurements for that high-accumulation condition.

Replicate variation in WT PHA content was high under severe nutrient limitation, with coefficient of variation 44% at 33% phosphorus and 37% at 8% nitrogen. Within these groups, final optical density varied by only 7–9% (Data 13 in Data Set S1). The variability therefore also reflects differences in PHA accumulated per unit biomass, reflecting that the timing of nutrient depletion varied between replicates due to a fixed harvest time. The high within-group variability in the WT 33% phosphorus condition (CV ∼44%) should be taken into account when interpreting the reported 3.7-fold difference between Δ*phaZ1* Δ*phaZ2* and WT at this condition; the fold-change is statistically supported but carries greater uncertainty than comparisons made under less severe nutrient limitation.

### PHA accumulation increased via a two-stage cultivation

To improve PHA accumulation while maintaining biomass formation, we tested a two-stage growth strategy. In Stage 1, cells were grown under nutrient-replete conditions to support biomass accumulation until stationary phase. In Stage 2, cultures were transferred to nitrogen (8%) and phosphorus (33%) limited medium containing excess carbon to promote PHA storage. In the one-stage control, cells were transferred from nutrient-replete medium into fresh nutrient-replete medium (Fig. 5A).

**Figure 5.**
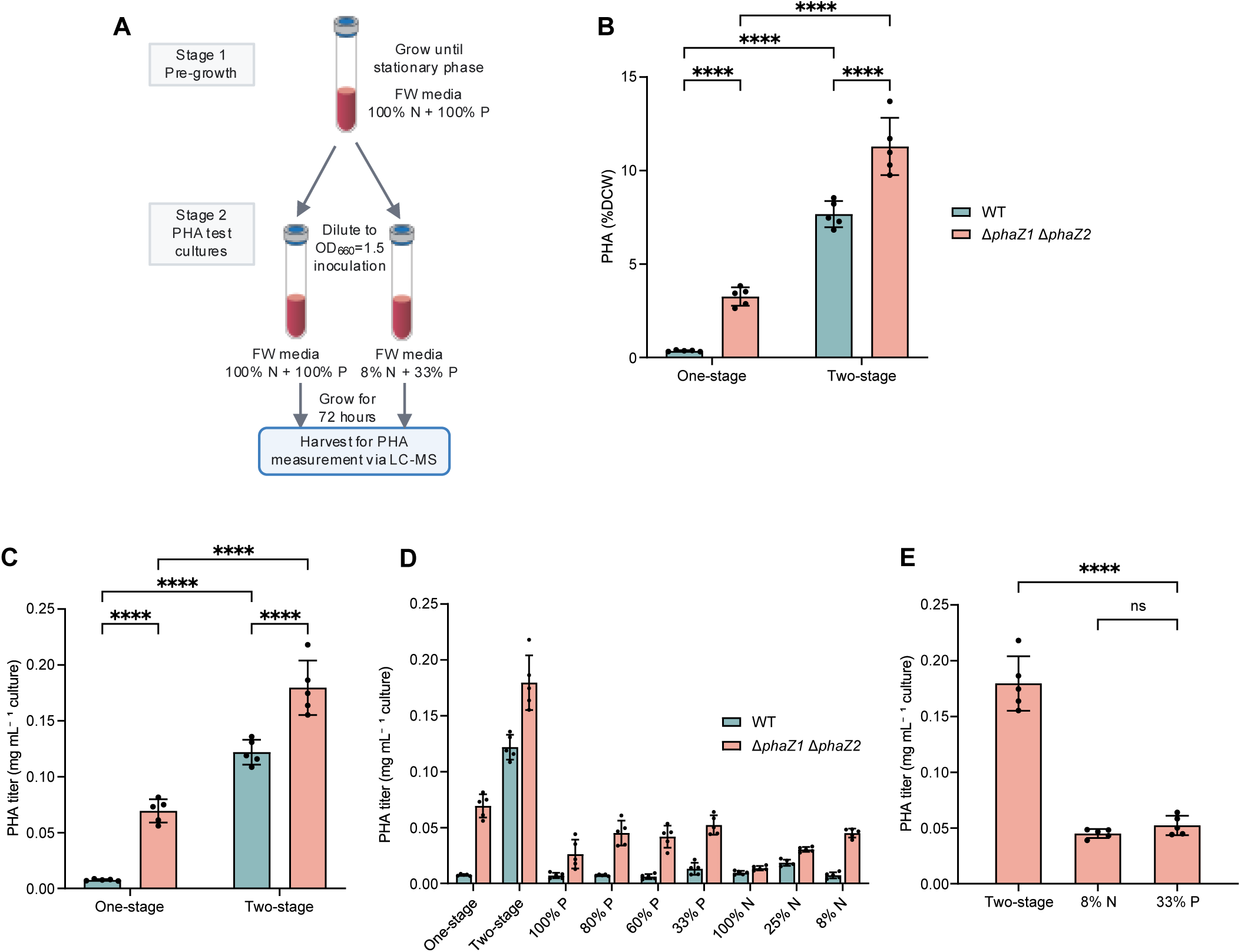
Two-stage cultivation increases intracellular PHA content and volumetric PHA titer. (A) Two-stage cultivation workflow. Wild type and *ΔphaZ1 ΔphaZ2* were first grown to stationary phase in nutrient-replete freshwater medium, diluted to OD₆₆₀ = 1.5, and transferred either to fresh nutrient-replete medium (one-stage control; 100% nitrogen and 100% phosphorus) or to storage medium containing 8% nitrogen and 33% phosphorus simultaneously (two-stage cultivation); this combined nitrogen and phosphorus limitation differs from the single-nutrient experiments in Fig. 4, where nitrogen and phosphorus were varied independently. Cultures were harvested after 72 h. (B) PHA content as a percentage of dry cell weight. (C) Volumetric PHA titer (mg PHA mL⁻¹ culture). (D) Volumetric PHA titer for wild type and Δ*phaZ1* Δ*phaZ2* across all cultivation conditions tested in this study, including the phosphorus- and nitrogen-limitation series shown in Fig. 4. (E) Volumetric PHA titer of Δ*phaZ1* Δ*phaZ2* under two-stage cultivation compared with the two single-stage nutrient-limited conditions that gave the highest titer. Data in B and C were analyzed by ordinary two-way ANOVA with genotype and cultivation strategy as fixed factors, including their interaction, followed by Šídák-adjusted comparisons between genotypes within each cultivation strategy and between cultivation strategies within each genotype. Comparisons in panels B and C are within-experiment (all cultures from the same inoculum and harvest batch); the statistical test therefore captures true cultivation-strategy effects rather than cross-experiment variation. In E, titer under two-stage cultivation was compared with titer under 8% nitrogen and 33% phosphorus limitation by Brown-Forsythe and Welch ANOVA with Dunnett’s T3 multiple-comparisons test, using two-stage cultivation as the control group; because these conditions were assessed in separate experiments, this comparison is descriptive and differences may reflect variation in medium composition, inoculum density, and harvest time in addition to cultivation strategy. No statistical comparisons are shown in D. Points represent biological replicates (n = 5); bars show the mean with standard deviation. ns, not significant; ****P < 0.0001. †FTIR-estimated PHA values exceeding the calibration maximum (3.89% DCW) are outside the validated calibration range; absolute values for these data points should be interpreted as estimates only.

Two-stage cultivation substantially increased PHA content, approximately 21.5-fold in WT and 3.5-fold in *ΔphaZ1 ΔphaZ2* relative to their one-stage controls (*P* < 0.0001 for both), and the double mutant accumulated approximately 1.5-fold more PHA than WT under two-stage cultivation (Fig. 5B). Two-way ANOVA identified significant effects of cultivation strategy and genotype (both *P* < 0.0001), accounting for 81.6% and 14.8% of the total variation respectively, with no significant interaction (*P* = 0.374).

Volumetric PHA titer followed the same pattern, increasing approximately 16.1-fold in WT and 2.6-fold in Δ*phaZ1* Δ*phaZ2* relative to their one-stage controls (*P* < 0.0001 for both), with the double mutant reaching a titer approximately 1.5-fold higher than WT under two-stage cultivation (*P* < 0.0001) (Fig. 5C). Cultures reached OD□□□ of approximately 2.6 under two-stage cultivation, compared with approximately 0.33 under single-stage 33% phosphorus limitation, indicating that biomass formation was largely preserved.

Comparing volumetric titer across all cultivation conditions tested, two-stage cultivation produced the highest titer in both strains (Fig. 5D). In the double mutant, titer under two-stage cultivation was approximately 3.4-fold higher than under 33% phosphorus limitation and 4.0-fold higher than under 8% nitrogen limitation, despite lower PHA content per unit biomass under two-stage conditions.

### FTIR spectroscopy provided a higher-throughput readout of PHA accumulation

Our previous work used LC–MS to quantify PHA in small culture volumes (5,14,15). However, that methodology is labor-intensive, limiting sample throughput. Here, we evaluated FTIR spectroscopy as a faster method to monitor PHA accumulation in TIE-1 by detecting the intact polymer. Representative FTIR spectra showed clear differences between a strong PHA-accumulating sample and a weak PHA-accumulating sample in the PHA-associated carbonyl region (Fig. S2). To quantify this signal, we focused on the PHA carbonyl peak at approximately 1700–1750 cm⁻¹ and normalized it to the biomass-associated amide I and amide II regions at approximately 1600–1700 cm⁻¹ and 1500–1580 cm⁻¹, respectively (Fig. S2). This normalization accounted for differences in biomass loading between samples.

The FTIR workflow included cell harvesting, washing, lyophilization, FTIR measurement, baseline correction, peak integration, and calibration against LC–MS-based PHA measurements (Fig. 6A). To build the calibration model, late stationary-phase samples were analyzed by both LC–MS and FTIR. LC–MS-measured PHA content was used as the quantitative reference for PHA % DCW readouts, while the FTIR carbonyl/amide peak-area ratio was used as the predictor. The FTIR carbonyl/amide peak-area ratio strongly correlated with LC–MS-measured PHA % (DCW). Linear regression produced the calibration equation:

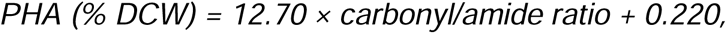

with an R² value of 0.928, and *n*=15, with a final-model RMSE of 0.314% DCW and MAE of 0.240% DCW (Fig. 6B). To evaluate predictive performance, leave-one-out cross-validation was performed. The cross-validated model retained similar accuracy, with an RMSE of 0.354% DCW, an MAE of 0.273% DCW, and an R² of 0.908. The resulting linear calibration model was then applied to exponential- and early stationary-phase samples that were analyzed by FTIR only. This model was calibrated over PHA contents between 0.57–3.89% DCW using 15 late-stationary-phase samples analyzed by both FTIR and LC–MS, and was applied only within that range to predict PHA content of the samples except for 2 measurements. These two values (4.03 and 4.49% DCW, both at 77 h under 25% nitrogen) exceed the calibration maximum of 3.89% DCW and are extrapolations; they should be interpreted as estimates only.

**Figure 6.**
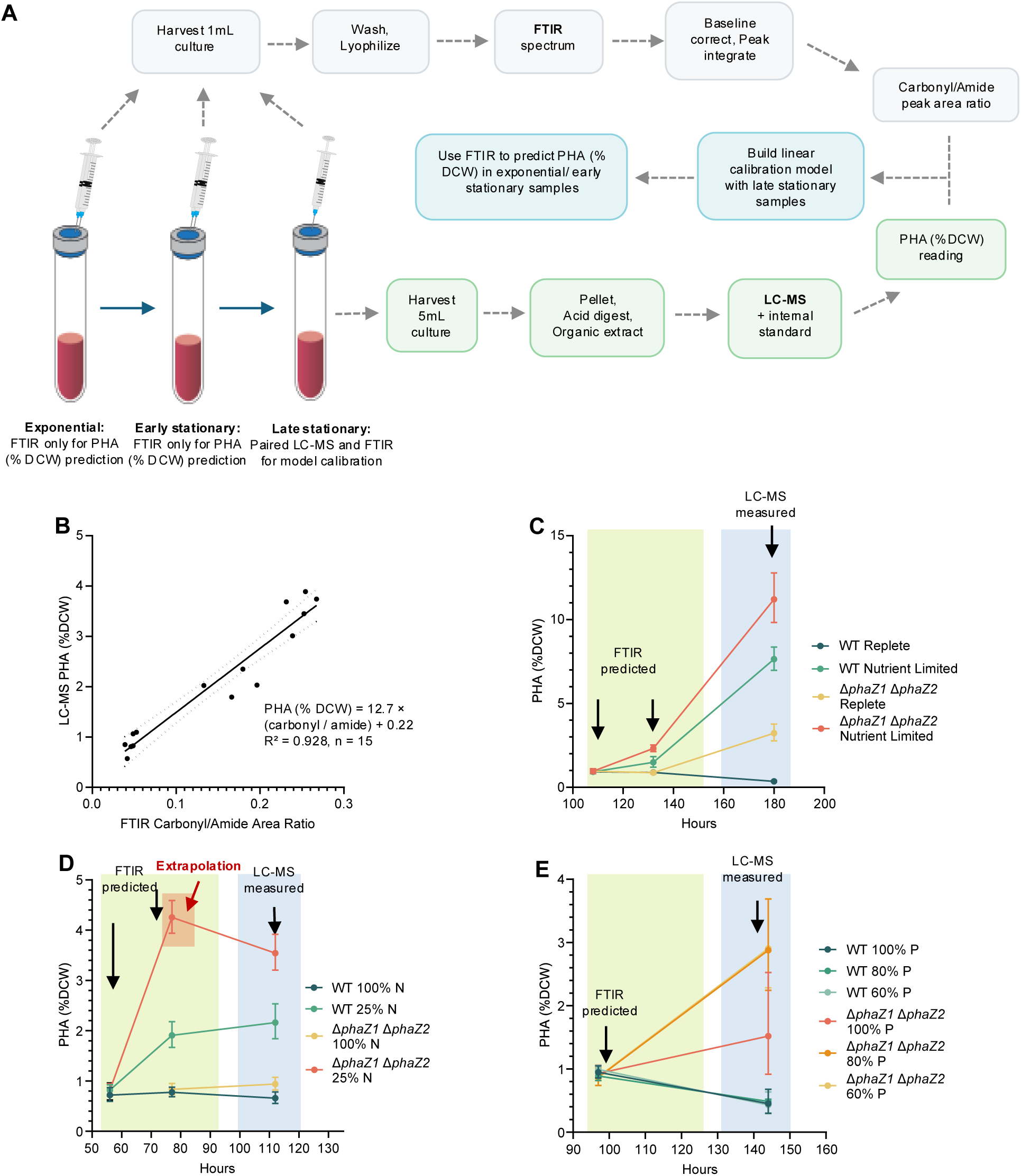
LC–MS-calibrated FTIR analysis of PHA accumulation. (A) Experimental workflow. Washed, lyophilized cells were analyzed by FTIR, and late-stationary-phase samples were analyzed in parallel by LC–MS to calibrate the carbonyl-to-amide ratio against PHA content. The resulting model was applied to FTIR measurements from exponential- and early-stationary-phase samples. (B) Linear relationship between the FTIR carbonyl-to-amide peak-area ratio and LC–MS-measured PHA content in the calibration samples (n = 15); the line denotes the fitted linear regression (PHA % DCW = 12.70 × ratio + 0.220; R² = 0.928), and the shaded band is the 95% prediction interval for the curve, which spans approximately ±0.75% DCW across the calibrated range. (C–E) FTIR-predicted PHA content at earlier sampling times and LC–MS-measured PHA content at the late-stationary endpoint during one-stage and two-stage cultivation, that 108 h is right before the split and transfer to the storage medium (C), under 100% and 25% nitrogen availability (D), and under 100%, 80%, and 60% phosphorus availability (E). Values are presented as the mean with standard deviation. FTIR estimates are intended for calibrated, semi-quantitative comparisons; LC–MS was used as the reference method for absolute PHA quantification. No statistical comparisons are shown in panels C–E; the time-course data are presented as descriptive summaries of FTIR-predicted PHA accumulation trends across strains and conditions.

When applied across growth stages and cultivation conditions, FTIR-estimated PHA values captured the major trends observed by LC–MS. In the two-stage cultivation experiment, FTIR predictions showed increased PHA accumulation after transfer to storage conditions, with the *ΔphaZ1 ΔphaZ2* mutant reaching the highest predicted PHA content (Fig. 6C). Similarly, under nitrogen limitation, FTIR-predicted PHA accumulation increased over time and was higher in the *ΔphaZ1 ΔphaZ2* mutant than in WT under nitrogen-limited conditions (Fig. 6D). Under phosphorus limitation, FTIR predictions also showed increased PHA retention in the double mutant under stronger phosphorus limitation, although higher replicate variation was observed in the limited conditions (Fig. 6E). Together, these results support FTIR as a higher-throughput method for screening PHA accumulation titers across TIE-1 strains and growth conditions.

## Discussion

### PhaZ1 and PhaZ2 are evolutionarily distinct but functionally redundant PHA depolymerases

Rather than being controlled by a single depolymerase, PHA turnover in TIE-1 appears to involve two enzymes with overlapping functions. PhaZ1 and PhaZ2 fall into separate clades on the protein tree, indicating that they are divergent evolutionarily, and their domain architectures differ accordingly. PhaZ1 groups with putative intracellular short-chain-length PHA depolymerases and contains a DepA/PHA depolymerase-like domain, consistent with a role in intracellular PHA mobilization. PhaZ2 groups with putative extracellular short-chain-length PHA depolymerases and contains a CE1/LpqC esterase-like hydrolase domain (24–26), yet the signal peptide upstream of its coding sequence is separated from it by an in-frame stop codon, so the expressed protein is not predicted to be secreted. The genetic data nonetheless indicate that *phaZ2* encodes a functional product, since its deletion increases PHA retention in the Δ*phaZ1* background, and expression in trans shortens lag time. A disrupted export signal immediately upstream of a functional depolymerase can be explained by loss of an ancestral signal peptide, with the catalytic domain retained and the product remaining cytoplasmic. Hence, its clade placement might reflect similarity to secreted enzymes, with the absence of a predicted signal on the expressed protein. This model also accounts for the behavior of PhaZ2 in trans: a cytoplasmic enzyme would not be separated from the polymer by a membrane, and its activity would instead be limited by access to the granule surface. Direct determination of PhaZ2 localization and transcript structure at this locus would test this model.

Under both non-nitrogen-fixing and nitrogen-fixing photoheterotrophic growth with butyrate, deletion of *phaZ1*, *phaZ2*, or both did not significantly alter lag time or final optical density, but all three deletion strains showed longer doubling times than wild type, with the largest effect in the double mutant. The same growth phenotype was observed in a *ΔphaZ1* strain in our previous work (5). Because PHA metabolism is dynamic rather than purely a storage endpoint, with cells both synthesizing and remobilizing polymer according to nutrient availability and metabolic demand (11–13), the longer doubling times are consistent with a role for PHA turnover in carbon use during growth, although complementation did not restore doubling time.

PHA retention followed a distinct genetic pattern. Deletion of either gene alone did not substantially increase PHA content under either condition, whereas the double mutant accumulated approximately 2.3-fold more PHA than wild type under non-nitrogen-fixing conditions and approximately 1.7-fold more under nitrogen-fixing conditions. The double-mutant phenotype therefore exceeds what the individual single-mutant effects would predict, indicating that loss of one depolymerase is buffered by the other.

Studies in other bacterial models have revealed similar genetic patterns in PHA turnover. In *Ralstonia eutropha* H16, now *Cupriavidus necator*, multiple intracellular PhaZ enzymes contribute to PHA turnover. In this system, PhaZ1 and PhaZ2 have overlapping but different functions, where the loss of both enzymes produces a stronger PHA-retention phenotype than loss of either enzyme alone. Further studies also show that different PhaZs in *R. eutropha* contribute to PHA mobilization, oligomer degradation, and granule organization, supporting the idea that PHA turnover may require multiple depolymerase activities with overlapping but distinct functions (27–31). A comparable double depolymerase deletion was recently reported in Marinibacterium sp. CCB-SX1, where simultaneous knockout of two *phaZ* genes substantially increased PHA accumulation; notably, in that organism individual deletions produced detectable phenotypes, whereas in TIE-1 neither single deletion substantially elevated PHA content under the conditions tested.

The predicted domain architecture of TIE-1 PhaZ2 is not a direct match to the *R. eutropha* PhaZ model, however. In *R. eutropha*, PhaZ2 is an intracellular 3-hydroxybutyrate-oligomer hydrolase acting alongside other cytoplasmic depolymerases, whereas TIE-1 PhaZ2 groups with the extracellular clade and carries a CE1/LpqC esterase-like domain. By sequence, it is closer to the depolymerase described in *Rhodospirillum rubrum*, where an enzyme originally annotated as intracellular was shown to carry a functional signal peptide and localize to the periplasm; the authors proposed that a second PhaZ-like gene lacking a signal peptide encodes the true intracellular enzyme (27,32). The second enzyme was subsequently purified and shown to hydrolyze native PHA granules with high activity (31). The parallel is incomplete, however: the *R. rubrum* enzyme retains a functional signal peptide and localizes to the periplasm, whereas the signal peptide upstream of TIE-1 *phaZ2* is interrupted. Both cases nonetheless illustrate that clade assignment and annotation do not determine localization. Because the localization and biochemical activity of TIE-1 PhaZ2 have not been determined experimentally, its mechanism and site of action remain open questions.

PHA accumulation was higher under nitrogen-fixing than non-nitrogen-fixing photoheterotrophic growth, consistent with recent work from our group (5,14), but differing from our previous observations under photoferrotrophic and photoelectrotrophic growth. Nitrogen fixation therefore does not have a uniform effect on PHA accumulation in TIE-1; its impact depends on metabolic context. Under butyrate-fed photoheterotrophic conditions it may shift carbon and reducing power toward storage, whereas under autotrophic conditions its energetic cost may instead reduce net accumulation (18).

### PhaZ1 and PhaZ2 promote PHA turnover under nutrient-limited storage conditions

Because PHA storage is commonly stimulated when carbon is available in excess relative to growth-limiting nutrients, we tested growth and PHA storage under nitrogen and phosphorus limitation. Under nutrient-replete conditions, PHA accumulation remained low. Although *ΔphaZ1 ΔphaZ2* showed a modest increase in PHA content relative to WT under 100% nitrogen and 100%, 80%, and 60% phosphorus conditions, these differences were not statistically significant after two-way ANOVA with multiple-comparison correction. By contrast, significant genotype-by-nutrient interactions were detected in both the nitrogen- and phosphorus-limitation experiments, indicating that the effect of phaZ1 phaZ2 deletion depended on nutrient availability. The strongest PHA-retention phenotype was observed under severe but still growth-permissive nutrient limitation, at 8% nitrogen and 33% phosphorus. These severe-limitation conditions also showed greater replicate-to-replicate variation, likely because reduced biomass and increased PHA accumulation made PHA % DCW estimates more sensitive to differences in growth, sampling, and extraction. Nevertheless, the significant interactions and condition-specific comparisons support the conclusion that the double-mutant phenotype became strongest as nitrogen or phosphorus availability decreased.

Severe nutrient limitations likely restrict biomass formation while leaving carbon source from butyrate available, promoting carbon storage. It was previously shown that both nitrogen and phosphorus limitation can enhance PHA accumulation in activated sludge (33). In this context, WT cells may continue to remobilize PHA despite at a reduced rate, whereas the *ΔphaZ1 ΔphaZ2* mutant is stunted in stored PHA turnover. However, because PHA synthesis and degradation were not measured kinetically, the increased endpoint PHA content cannot distinguish reduced remobilization from additional synthesis. In addition, when limitations become too strong, such as under 0% nitrogen with argon headspace or 5% phosphorus, growth is too limited to generate sufficient biomass for PHA analysis. Therefore, the strongest PHA-retention phenotype occurs under severe but still growth-permissive nutrient limitation, indicating that PhaZ1 and PhaZ2 contribute to PHA turnover most clearly when TIE-1 metabolism is shifted toward storage.

### Complementation attributes the PHA phenotype to the deleted genes

Expression of *phaZ1* and *phaZ2* in trans reduced PHA content in the *ΔphaZ1 ΔphaZ2* strain toward wild-type levels, indicating that the PHA retention phenotype is attributable to loss of these genes. Restoration was incomplete, which might be caused by expression from a plasmid-borne inducible promoter that reproduces neither the level, the timing, nor the regulation of native expression.

Because the complementation strains were grown at 25% nitrogen availability with 5 mM butyrate rather than the nutrient-replete, 10 mM butyrate conditions used for the deletion panel (Fig. 2), absolute PHA contents are not directly comparable between the two experiments. However, the larger wild-type-to-double-mutant difference observed here (3.1-fold versus 2.4-fold) is consistent with the amplification of this phenotype under nitrogen limitation.

### PhaZ1 and PhaZ2 exhibited different activity in trans

The two enzymes behaved differently in trans across three different readouts. Neither single mutant had a PHA phenotype to restore, consistent with the deletion panel (Fig. 2), so expression in these backgrounds tested depolymerase dosage rather than restoration. Expression of *phaZ1* lowered PHA content from 4.1% to 1.7% DCW (*P =* 0.0413), approximately half the wild-type level, shortened lag time (*P =* 0.0114), and increased doubling time (*P =* 0.0344). Expression of *phaZ2* shortened lag to a comparable degree (10.7 versus 9.8 h; *P =* 0.0054) but changed neither PHA content, 4.9% versus 4.8% DCW (*P* > 0.999) nor doubling time (*P =* 0.2073), despite being driven from the same vector under the same growth and induction conditions. PhaZ2 activity in vivo therefore does not appear to be limited by enzyme abundance, instead potentially by localization or access to the polymer. This is consistent with its predicted extracellular clade placement and with observations in *R. eutropha*, where the activity of one depolymerase paralog increased up to 21-fold when the proteinaceous granule surface layer was removed (31). In *R. rubrum*, the activator of PHA degradation was identified as ApdA, which is a phasin that stimulates hydrolysis of PHA granules without proteolytically modifying the surface layer (34). Phasins have been identified in *R. palustris*, where RPA0089 and RPA3770 appear to be the granule-associated proteins and RPA0089 expression increases progressively during nitrogen starvation, suggesting that granule surface composition may also influence depolymerase access (35,36).

Three non-exclusive hypotheses may explain the failure of *phaZ2* to reduce intracellular PHA when expressed in *trans*. First, PhaZ2 may require a co-factor or activator protein analogous to ApdA in *R. rubrum* (34): if PhaZ2 activity depends on an interaction partner not provided by the vector context, enzyme abundance alone would not restore depolymerase function. Second, PhaZ2 may fail to reach the granule surface when produced ectopically; if its native context includes a signal-peptide-independent targeting mechanism that is disrupted by vector-borne expression, the enzyme could accumulate in the cytoplasm or periplasm without accessing substrate. Third, the Rpal_0568 gene product may lack *in vivo* depolymerase activity entirely under tested conditions despite its domain similarity to characterized esterases; in that case, the complementation result would correctly reflect a null activity. Distinguishing among these hypotheses will require direct localization studies, *in vitro* activity assays, and identification of potential protein interaction partners. Biochemical confirmation of PhaZ1 and PhaZ2 depolymerase activity through *in vitro* assays with purified protein and native PHA granules would be required to formally establish their enzymatic function, independent of the genetic evidence.

Placement of PhaZ2 in the putative extracellular clade reflects sequence and domain similarity to characterized extracellular depolymerases rather than demonstrated secretion. SignalP 6.0 analysis of the annotated Rpal_0568 product (PhaZ2) predicted no signal peptide, with a likelihood of 1.00 for the no-signal-peptide class (Fig. S3). A cytoplasmic or granule-associated localization of PhaZ2 is therefore compatible with the clade assignment, and it would be limited, if at all, by access to the granule surface as documented in *R. rubrum* (31), although localization has not been determined experimentally. The contribution of *phaZ2* to intracellular PHA retention, inferred from the Δ*phaZ1* Δ*phaZ2* phenotype, presents a mechanistic question that the SignalP prediction alone does not resolve: if PhaZ2 is functionally homologous to extracellular depolymerases yet lacks a detectable Sec/SPI signal peptide, how does it access the PHA granule? A useful precedent is the *Rhodospirillum rubrum* PHA depolymerase PhaZa1, initially annotated as intracellular but subsequently shown to carry a functional signal peptide and localize to the periplasm; its activity on native granules was demonstrably enhanced by removal of the proteinaceous granule surface layer (Handrick et al., 2004, J. Bacteriol. 186:7243). For TIE-1 PhaZ2, the in-frame stop codon interrupting the upstream signal-peptide ORF (Fig. 1C) prevents canonical Sec/SPI export, making cytoplasmic or granule-associated localization the most parsimonious inference; nevertheless, other secretion pathways not detected by SignalP 6.0 cannot be formally excluded. Direct determination of PhaZ2 localization, by subcellular fractionation or fluorescent-protein fusion, would clarify whether it acts on granules from the cytoplasm or via an unrecognized export route.

Because the two open reading frames in the tandem construct were not assayed separately, the contribution of each to the partial rescue of the double mutant cannot be resolved, and a predominantly *phaZ1*-driven rescue cannot be excluded. The double-mutant comparison nonetheless remains a genuine test of restoration, since that background, unlike either single mutant, shows a strong PHA phenotype.

### Lag time and growth rate respond to depolymerase overexpression in opposite directions

Lag time and exponential growth rate responded to depolymerase expression in opposite directions. Both single complements shortened lag while *phaZ1*-containing constructs lengthened doubling time, and final optical density was unaffected throughout. Expression from the vector-borne promoter was induced at inoculation, so depolymerase was present during the lag phase as well as during exponential growth. Hence, a plausible interpretation is that mobilization of PHA from rich media inoculum supports the transition out of lag. However, simultaneous synthesis and degradation of PHA consumes ATP and NADPH once balanced growth is underway (37).

Depolymerase activity therefore affects growth rate in both directions: deletion slows growth, and overexpression slows it as well. Constant and unregulated expression from the plasmid, rather than failed complementation, therefore accounts for the longer doubling times, and this does not undermine the attribution of the PHA phenotype to loss of *phaZ1* and *phaZ2*. Plasmid maintenance and antibiotic selection also impose a cost in this organism: a growth defect in TIE-1 complementation strains carrying pBBR1MCS-5 has previously been attributed to the combined effect of the plasmid and gentamicin (7). Because all strains in this experiment carried pSRKGm and were cultured with gentamicin, this cost applies equally across comparisons, and the divergent growth outcomes are attributable to the genes expressed rather than to plasmid maintenance. A comparable cost from depolymerase expression specifically has been reported in *R. eutropha*, where constitutive expression of PhaZd1 or PhaZd2 from a pBBR1-derived vector prevented PHA accumulation and PhaZd2-expressing cells grew more slowly than wild type (31).

### Two-stage cultivation method maintained biomass while increasing PHA yield

Although nitrogen- and phosphorus-limited conditions increased PHA content in the *ΔphaZ1 ΔphaZ2* double mutant relative to WT under nutrient-replete conditions, biomass accumulation was lower due to limited nutrients. Two-stage cultivation has been widely used in bioplastic production to resolve the tradeoff between biomass formation and polymer storage. In mixed-culture PHA systems, biomass is first generated or enriched under growth-supporting conditions and then transferred to a production stage with excess carbon and limited nutrients to minimize further growth while maximizing intracellular polymer content (10,38,39).

Our two-stage cultivation strategy allowed cells to first accumulate biomass under nutrient-replete conditions and then shift toward PHA accumulation when nitrogen and phosphorus became limiting. This strongly increased PHA content in both WT and *ΔphaZ*1 *ΔphaZ*2, with the double mutant maintaining higher accumulation across both strategies. The significant effects of cultivation strategy and genotype, without significant interaction, indicate that two-stage cultivation and depolymerase deletion contributed additively rather than synergistically. This differs from the nutrient-limitation experiments, in which significant genotype-by-nutrient interactions showed that the double-mutant phenotype depended strongly on nitrogen or phosphorus availability.

Two-stage cultivation also increased volumetric PHA titer, approximately 16.1-fold in WT and 2.6-fold in *ΔphaZ1 ΔphaZ2* relative to their one-stage controls. Meanwhile, the double mutant reached a titer approximately 23.6-fold higher than WT under one-stage cultivation. Notably, single-stage phosphorus limitation produced more than twice the PHA content per unit biomass (26.3% versus 11.3% DCW) yet approximately 3.4-fold lower volumetric titer, because biomass formation was severely restricted under that condition. PHA content and volumetric productivity are therefore not equivalent metrics, and maximizing cellular PHA fraction alone does not maximize yield.

Simultaneous deletion of *phaZ1* and *phaZ2* improved PHA retention most clearly when carbon remained available and nitrogen or phosphorus were limited. Because substrate consumption and production rates were not quantified, however, these results do not establish improved substrate yield or volumetric productivity. Based on the 72 h Stage 2 incubation period, the 0.180 mg mL ¹ titer in the double mutant corresponds to a volumetric productivity of approximately 0.0025 mg mL□¹□h□¹, which is at the low end of reported PNSB systems and highlights the need for process intensification. The 0.180 mg mL⁻¹ volumetric titer achieved here is approximately 3.4-fold higher than the highest previously reported for TIE-1 (0.053 mg mL⁻¹) (5) but remains below the 0.377 mg mL⁻¹ reported for *Rhodopseudomonas* sp. S16-VOGS3 in a fed-batch outdoor photobioreactor (48), indicating that further carbon loading, process optimization, and strain engineering will be needed to approach industrially relevant yields. Among orthogonal strategies tested in TIE-1, overexpression of RuBisCO form I and II genes increased PHA accumulation under both heterotrophic and photoautotrophic conditions (5); combining depolymerase deletion with enhanced carbon fixation may therefore offer an additive path toward closing this gap.

### LC–MS-calibrated FTIR enables rapid screening of PHA accumulation

FTIR-based methods have previously been developed as rapid approaches for PHA quantification (40), particularly in mixed-culture and wastewater-based production systems. Arcos-Hernandez *et al.* showed that infrared spectra could be calibrated against PHA content to rapidly quantify intracellular PHA in mixed cultures, reducing analytical time compared with conventional GC-based methods. More recently, Deng et al. optimized FTIR spectroscopy for quantifying PHA production in municipal wastewater sludge and further supported the use of FTIR as a practical analytical method for microbial mixed-culture PHA production (15–17).

Similarly, our FTIR-based method provides a high throughput approach to monitor PHA accumulation in TIE-1. By pairing FTIR measurements with LC–MS measurements from the same late stationary-phase samples, we built a calibration model linking the FTIR carbonyl/amide peak-area ratio to quantitatively measured PHA content. This approach uses LC–MS as the quantitative reference method while allowing FTIR to serve as a faster readout that requires less biomass and hence can be applied across different growth stages of samples, revealing temporal resolution in TIE-1’s PHA production. Importantly, FTIR-predicted PHA values reproduced the major biological trends observed by LC–MS, including increased PHA retention in the Δ*phaZ1* Δ*phaZ2* mutant under nutrient-limited and two-stage storage conditions.

Because FTIR estimates PHA indirectly from spectral peak area ratios, this method is best used as a semi-quantitative screening tool rather than a complete replacement for LC–MS. Variation in biomass loading, spectral acquisition, baseline correction, and peak integration can affect the carbonyl/amide ratio, especially for samples that have low PHA accumulation. Therefore, LC–MS remains important for absolute PHA quantification and model calibration. However, once calibrated, FTIR provides a useful way to compare relative PHA trends across strains, growth stages, and nutrient conditions, making it valuable for screening PHA accumulation phenotypes in TIE-1, consistent with the view that spectroscopic methods are best positioned as rapid screens calibrated against chromatographic reference methods (15).

The strong correlation between the FTIR peak-area ratio and LC–MS-measured PHA content further supports the validity of the model because the two methods measure PHA through different chemical readouts. LC–MS quantifies PHA indirectly after acid digestion, in which PHA is converted to crotonic acid-derived products used as a proxy for polymer abundance. In contrast, FTIR measures intact dried biomass and detects the PHA-associated carbonyl absorbance from the polymer itself. Consistent with this interpretation, low-storage conditions showed weak carbonyl peaks, whereas nutrient-limited, high-storage conditions showed stronger carbonyl absorbance. The FTIR carbonyl/amide ratio therefore tracks differences in PHA accumulation and is unlikely to reflect variation in biomass loading or spectral processing alone. This cross-method agreement strengthens the use of FTIR as a rapid screening method for PHA accumulation in TIE-1.

The small difference between the final-model and cross-validated prediction errors indicates that the calibration was not strongly overfit to the available dataset. A cross-validated error of approximately 0.35% DCW is sufficiently small for comparing relative PHA accumulation among strains and nutrient conditions.

Future work should first establish the molecular basis of the double-mutant phenotype with RT-qPCR. Time-resolved measurements of PHA synthesis and remobilization would further distinguish reduced degradation from changes in polymer synthesis or biomass allocation. For nutrient-limitation and two-stage experiments, 5 mM butyrate was used to maintain cultures within a growth range suitable for comparing strain performance and nutrient responses. Because carbon availability can influence PHA accumulation, higher butyrate concentrations may be used in future optimization experiments to increase PHA content and total PHA production. Finally, the FTIR model should be evaluated prospectively using independent biological batches spanning different genotypes, growth phases, nutrient conditions and PHA concentrations. These studies would define the model’s domain of applicability and determine whether it can support reliable high-throughput screening during subsequent strain and process optimization.

The calibration set was also distributed bimodally, so the R² should not be read as evidence of within-condition precision. Of the samples to which the model was applied, 6 out of 94 samples fell in the unsampled interval between clusters. Finally, the environmental benefit of PHA produced by this route was not assessed; determining whether it offers advantage in environmental impact over petroleum-derived plastics would require life-cycle assessment of the full cultivation and recovery process. PHA monomer composition was not determined in this study; PHB is assumed based on prior characterization of TIE-1 polymer (14), but GC-MS compositional analysis would be required to confirm monomer identity under the nutrient conditions used here.

## Materials and Methods

### Bacterial Strains, Media, Growth Conditions

*Escherichia coli* strains were grown in lysogeny broth (LB) at 37°C. For chemoheterotrophic pre-growth of *Rhodopseudomonas palustris* TIE-1, strains were grown aerobically at 30°C in YPS-MOPS medium containing 3 g/L yeast extract, 3 g L⁻¹ peptone, 10 mM MOPS [3-(N-morpholino)propane sulfonic acid], and 10 mM succinate, adjusted to pH 7.2.

### Photoheterotrophic anaerobic growth of TIE-1

For PHA accumulation experiments, TIE-1 cultures were first grown in YPS-MOPS medium and then transferred into Freshwater Minimal Medium, containing NH_4_Cl, KH_2_PO4, CaCl_2_·2H_2_O, and MgSO_4_·7H_2_O, vitamins, and trace elements, supplemented with butyrate as the sole carbon source. All photoheterotrophic growth was performed anaerobically at 30°C under constant illumination.

### WT and PHA depolymerase mutants growth experiments under nitrogen-fixing and non-nitrogen-fixing conditions

For growth comparisons under nitrogen-fixing and non-nitrogen-fixing conditions, cultures were grown with 1 mM butyrate. For PHA-accumulation assays comparing nitrogen-fixing and non-nitrogen-fixing conditions, cultures were grown with 10 mM butyrate and collected at early stationary phase. For non-nitrogen-fixing conditions, freshwater medium was supplemented with NH₄Cl. For nitrogen-fixing conditions, NH₄Cl was excluded from the medium.

### Complementation growth and PHA accumulation

For complementation experiments, wild type and each deletion strain carrying either empty pSRKGm or the corresponding expression construct were grown photoheterotrophically in freshwater medium containing 5 mM butyrate at 25% nitrogen availability, with the headspace exchanged with argon. Gentamicin (800 µg mL⁻¹) was included to maintain the plasmid, and IPTG (10 mM) was added at inoculation to induce expression. All strains were cultured in parallel so that plasmid maintenance, selection, and induction were matched across comparisons. Growth was monitored by OD₆₆₀ and cultures were harvested at 48 hours after cultures reach early stationary for PHA quantification as described above. Expression from pSRKGm constructs was not independently verified by RT-qPCR or Western blot; the absence of a detectable PHA phenotype in ΔphaZ2/pphaZ2 could therefore reflect insufficient induction or poor protein accumulation in addition to the biological mechanisms discussed above.

### Nutrient limitation growth experiments

For nitrogen-limitation experiments, NH₄Cl concentration was varied while butyrate was held constant at 5 mM for both growth and PHA accumulation. Nutrient-replete freshwater medium contains 0.3 g L⁻¹ NH₄Cl (5.6 mM); nitrogen conditions are expressed throughout as a percentage of this concentration, giving 0.075 g L⁻¹ (1.4 mM) at 25%, 0.024 g L⁻¹ (0.45 mM) at 8%, and none at 0%. Similarly, replete medium contains 0.5 g L⁻¹ KH₂PO₄ (3.7 mM), giving 0.40 g L⁻¹ (2.9 mM) at 80%, 0.30 g L⁻¹ (2.2 mM) at 60%, 0.165 g L⁻¹ (1.2 mM) at 33%, and 0.025 g L⁻¹ (0.18 mM) at 5%. Reduced media were prepared by mixing replete freshwater medium with nitrogen- or phosphorus-free freshwater medium at the appropriate ratios. Media completely lacking NH₄Cl or KH₂PO₄ were included as negative controls. Nitrogen-limiting experiments were performed with argon headspace to minimize nitrogen fixation.

### Two-stage cultivation

For two-stage cultivation experiments, WT and Δ*phaZ1* Δ*phaZ2* strains were first grown photoheterotrophically in nutrient-replete Freshwater Minimal Medium supplemented with 5 mM butyrate as the sole carbon source. Cultures were grown anaerobically until stationary phase and then diluted to OD₆₆₀ = 1.5 into either fresh nutrient-replete medium or nutrient-limited storage medium.

For the one-stage control, cultures were transferred into nutrient-replete Freshwater Minimal Medium containing 100% nitrogen and 100% phosphorus. For the two-stage cultivation condition, cultures were transferred into storage medium containing 8% nitrogen and 33% phosphorus, with butyrate maintained at 5 mM. Cultures were incubated photoheterotrophically for an additional 72 h before harvest. Biomass was collected at the endpoint to determine PHA content as percentage of dry cell weight and volumetric PHA titer as mg PHA mL⁻¹ culture.

### Growth measurements

Growth was monitored by measuring optical density at 660 nm with Spectronic 200 (Thermo Fisher Scientific, USA). Growth curves were used to compare the effects of depolymerase deletion on lag phase, doubling time, and final culture density under non–nitrogen-fixing, nitrogen-fixing conditions, and different nutrient limitation conditions.

### Growth kinetics quantified by model-fitting

Growth kinetics were quantified by fitting a modified Gompertz model to OD₆₆₀ time-course data from each biological replicate. For each replicate, measurements were ordered by time, and observations with missing, zero, or negative OD₆₆₀ values were excluded. OD₆₆₀ values were normalized to the first valid measurement for that replicate, and the response variable was calculated as the natural logarithm of relative optical density:

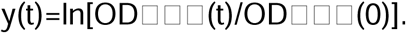

Growth curves were fit using the modified Gompertz equation:

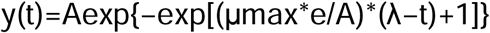

where *A* is the asymptotic increase in log OD₆₆₀, *µ*_max_ is the maximum specific growth rate (*h*^-1^), *λ* is the estimated lag time (*h*), and *t* is time (*h*). Models were fitted separately for each strain, condition, and biological replicate using bounded nonlinear least squares with the nlsLM function from the R package minpack.lm. The maximum number of iterations was set to 500.

Parameter bounds were set to 0.001–20 for *A*, 0.001–5 *h*^-1^for *µ*_max_, and 0 to the final sampling time for *λ* (23).

### Plasmid construction

All the plasmids used in this study are listed in Table 1. All the primers used are listed in Table S1. Deletion plasmids were constructed using the suicide vector pJQ200KS for markerless allelic exchange. For each target gene, approximately 1,000 bp upstream and 1,000 bp downstream homology arms flanking the coding sequence were PCR-amplified from *R. palustris* TIE-1 genomic DNA using primers listed in Table S1. The upstream and downstream fragments were designed to remove the majority of the target open reading frame while preserving the surrounding genomic context. PCR products were purified and assembled into linearized pJQ200KS by Gibson assembly. Assembled plasmids were transformed into *E. coli*, selected on gentamicin-containing plates, and screened by colony PCR and/or diagnostic restriction digestion. Correct plasmid construction was confirmed by Sanger sequencing across the assembled homology-arm junctions. Verified deletion plasmids were introduced into *E. coli* ABB9 for conjugation into *R. palustris* TIE-1.

For the construction of complementation plasmids, the coding sequences of *phaZ1* (*Rpal_0578*) and *phaZ2* (*Rpal_0568*) were amplified from wild-type *R. palustris* TIE-1 genomic DNA using primers carrying 5′ overhangs homologous to the pSRKGm multiple cloning site (Table S1). The broad-host-range expression vector pSRKGm was linearized by digestion with NdeI, and inserts were joined to the linearized vector by Gibson assembly, placing each gene under the control of the vector-borne IPTG-inducible *lac* promoter. For the tandem construct, phaZ1 and phaZ2 were assembled into the same site separated by a 20-bp linker (5′-ACTGTAGAGGAAATATCAGA-3′) containing a ribosome binding site (AGGAAA) positioned 7 nucleotides upstream of the phaZ2 start codon, placing both genes under the control of the vector-borne lac promoter as a bicistronic transcript. The native *phaZ1* stop codon was retained. Assembly reactions were transformed into chemically competent *E. coli* NEB 10-beta (New England Biolabs) and selected on LB agar containing gentamicin (100 µg mL□¹). Plasmids were verified by colony PCR and whole-plasmid sequencing before conjugative transfer into the corresponding TIE-1 deletion strains.

### Plasmid and Marker-less Deletion Strain Construction

Marker-less deletion strains were constructed by homologous recombination. Approximately 1 kb of sequence upstream and downstream of Rpal_0568 was amplified and cloned into the suicide vector pJQ200KS, which replicates in E. coli but not in *R. palustris* TIE-1. The cloned deletion plasmid was conjugated into either wild-type TIE-1 or ΔRpal_0578 TIE-1 using an E. coli donor strain ABB9. ABB9 is auxotrophic for diaminopimelic acid (DAP), and hence during conjugation, ABB9 was maintained on medium supplemented with 200 µM DAP. After mating with TIE-1, cells were plated on medium lacking DAP that counterselected ABB9, which isolated integrated TIE-1 strains. Single-crossover TIE-1 strains were selected by 800 μg mL⁻¹ gentamicin due to TIE-1’s high resistance to gentamicin, and double-crossover recombinant strains were counter-selected by sucrose. Resulting candidate strains were screened by PCR, and the deletion strains were confirmed by genomic sequencing. Genomic DNA extracted was submitted to Plasmidsaurus. Bacterial genome sequencing was performed by Plasmidsaurus using Oxford Nanopore Technology. To rule out secondary mutations introduced during the two rounds of allelic exchange, whole-genome sequencing of the *ΔphaZ1 ΔphaZ2* double mutant was performed.

### PHA extraction and LC–MS analysis

Intracellular PHA was quantified by LC–MS as crotonic acid following heat-treated methanol/chloroform/sulfuric acid digestion, as described previously (5,14,15), using 5 mL of culture per sample. Crotonic acid was detected on an Agilent Technologies 6420 Triple Quadrupole LC–MS at *m/z* 87. Because this assay detects crotonic acid as the pyrolysis product of 3-hydroxybutyrate units, it is specific to PHB; the presence of other monomers such as 3-hydroxyvalerate would not be captured by this measurement and would result in underestimation of total PHA content if the polymer is a copolymer.

PHA standards were prepared from dry PHA powder (Sigma-Aldrich) and processed through the same workflow as the experimental samples. A standard series was prepared and analyzed within each LC–MS sequence, and samples were quantified against the curve generated in the same experiment; standards spanned 0–1,000 ppm in the measurement used for Fig. 2 and 0–100 ppm in three other sequences. Coefficients of determination ranged from 0.958 to 0.995. Limits of detection and quantification, calculated as 3.3σ/S and 10σ/S, ranged from 0.20 to 62.15 ppm and from 0.59 to 188.34 ppm across sequences. In the Fig. 2 measurement, where both limits are highest, sample concentrations spanned 161–1,457 ppm, all above the limit of detection and 53 of 54 above the limit of quantification. Calibration data, per-standard recoveries, and this sensitivity analysis are provided in the supplemental material.

### Derivation of the OD_₆₆₀_ to dry cell weight conversion factor for Rhodopseudomonas palustris *TIE-1*

PHA abundance was normalized to biomass using the OD₆₆₀ measured at sample collection. Dry cell weight was estimated from optical density using a conversion derived specifically for TIE-1, following the approach used previously in this laboratory by Tahina et al. (5). The conversion combines a cell enumeration relationship with a measured average cell volume and literature values for fresh-cell density and dry-mass fraction:

DCW = cell concentration × cell volume × fresh-cell density × dry-mass fraction

### Cell concentration

Cell number was calculated from OD₆₆₀ using the enumeration relationship established for TIE-1 (5):

cells L⁻¹ = 1000 × 9.761951 × 10 × OD₆₆₀ × e^0.3068986 = 1.3268 × 10¹² × OD₆₆₀

### Parameters

| **Quantity** | **Value** | **Source** |
|---|---|---|
| Average cell volume | 1.048 µm<sup>3</sup> cell<sup>−1</sup> | Measured in TIE-1 by FM 1-43FX membrane staining and confocal imaging of ~1,000 individual cells (5) |
| Fresh-cell density | 1.1 g cm<sup>−3</sup> | (41) |
| Dry-mass fraction | 0.4 | (42) |
| Volume conversion | 10<sup>−12</sup> cm<sup>3</sup> µm<sup>−3</sup> | — |

As derived, 1 OD₆₆₀ unit corresponds to 0.612 mg dry cell weight mL⁻¹. This TIE-1-derived value agrees to within 2% with the factor of 0.625 g DCW L⁻¹ OD₆₆₀⁻¹ reported previously for *R. palustris* (43). Reference (43) determined this conversion factor gravimetrically, independently of the cell-volume-based derivation used here; the agreement between the two independent approaches provides cross-validation for the conversion factor applied throughout this study. Total dry cell weight in each culture sample was calculated as OD₆₆₀ × 0.612 × culture volume. Volumetric PHA titer was calculated by dividing the PHA mass recovered from each sample by the original culture volume and is reported as mg PHA mL⁻¹ culture.

Limitations of this method include that cell volume was measured in TIE-1, but fresh-cell density and dry-mass fraction are literature values treated as constant across strains and growth conditions. Reported dry-mass fractions for bacterial cells range from approximately 31 to 57% (42). Because PHA granules displace cell water and are denser than cytoplasm, the dry-mass fraction may be underestimated in high-PHA samples, which would underestimate dry cell weight and correspondingly overestimate PHA as a percentage of dry cell weight. However, fold-differences in PHA content between genotypes within a condition, and volumetric PHA titer, are independent of this conversion.

### FTIR based PHA quantification calibrated by LC-MS

Fourier-transform infrared spectroscopy was used as a higher-throughput method to estimate PHA accumulation. For paired FTIR and LC–MS analysis, each culture sample was split into aliquots so that FTIR measurements could be calibrated against LC–MS-derived PHA content from the same biological sample. For FTIR sample preparation, 1 mL culture samples were pelleted by centrifugation at 15000× g for 5 min. Cell pellets were washed three times with 0.9% NaCl to remove residual medium. After the final wash, supernatants were removed completely, and pellets were dried by lyophilization. Dried biomass pellets were analyzed using a Thermo Scientific Nicolet iS20 FTIR spectrometer to obtain infrared spectra.

FTIR spectra were baseline corrected before peak integration. PHA-associated signal was quantified by integrating the ester carbonyl region between 1700–1750 cm⁻¹ after subtraction of a flat baseline set to the minimum absorbance within the band. Biomass-associated signal was quantified by integrating the amide region, including amide I between 1600–1700 cm⁻¹ and amide II between 1500–1580 cm⁻¹. The PHA carbonyl peak area was normalized to the combined amide I and amide II peak area to generate a carbonyl-to-amide ratio for each sample:

Carbonyl-to-amide ratio = carbonyl area / (amide I area + amide II area)

To convert FTIR signals into estimated PHA content, carbonyl-to-amide ratios were calibrated against paired LC–MS measurements of PHA content reported as percent dry cell weight (% DCW). A linear calibration model was fit using the equation:

PHA % DCW = m × carbonyl-to-amide ratio + b,

where m is the fitted slope and b is the intercept. Model performance was evaluated using R², root mean squared error (RMSE), and mean absolute error (MAE), with RMSE and MAE reported in units of % DCW. To estimate predictive performance on held-out samples, leave-one-out cross-validation was performed. In each iteration, one sample was excluded, the linear model was trained using the remaining samples, and PHA content was predicted for the excluded sample. Cross-validated RMSE, MAE, and R² were calculated from these held-out predictions. The calibration set spanned LC–MS-measured PHA contents of 0.57 to 3.89% DCW and carbonyl-to-amide ratios of 0.039 to 0.267. Two samples at 77 h under 25% nitrogen gave predictions of 4.03 and 4.49% DCW, above the calibration maximum of 3.89% DCW; these are extrapolations while the rest of the prediction data are within the range. The calibration set was distributed bimodally, with a low-PHA cluster (0.57–1.09% DCW) from nitrogen-replete samples and a high-PHA cluster (1.79–3.89% DCW) from nitrogen-limited samples; the interval between 1.09 and 1.79% DCW was not represented in the calibration data and represents an unvalidated interpolation zone.

### Phylogenetic analysis of predicted PHA depolymerases

Candidate PHA depolymerase protein sequences were analyzed to evaluate the evolutionary relationship of the *R. palustris* TIE-1 depolymerase candidates with bacterial PHB/PHA depolymerases (Table S2). A total of 13 amino acid sequences, including the TIE-1 candidate proteins and representative PHA depolymerase homologs, were aligned and imported into MEGA12 for phylogenetic analysis.

Evolutionary relationships were inferred using the Neighbor-Joining method. Evolutionary distances were calculated using the p-distance method and are reported as the number of amino acid differences per site. Branch support was evaluated using 1,000 bootstrap replicates, and bootstrap values are shown next to the corresponding branches. Positions with less than 95% site coverage were removed using the partial deletion option, resulting in a final dataset of 370 aligned amino acid positions. The PHA depolymerase from *Afipia massiliensis* was used as the outgroup to root the tree. The final tree was visualized in MEGA12 and used to compare the clustering of TIE-1 PhaZ candidates with annotated intracellular and extracellular PHA depolymerase homologs (44–47).

### Domain architecture prediction

Predicted domain architectures of PhaZ1 (Rpal_0578) and PhaZ2 (Rpal_0568) were determined by searching the amino acid sequences against the NCBI Conserved Domain Database (CDD) using default search parameters. Domain boundaries are reported as amino acid positions within each protein (24–26). Signal peptides were predicted from the annotated protein sequences using SignalP 6.0 with default parameters.

### Statistical Analysis

Statistical analyses were performed using GraphPad Prism version 10.6. Estimated lag time, doubling time, and final optical density fitted from growth curves from individual biological replicates were used for statistical comparisons. Growth parameters and PHA content were compared among WT, *ΔphaZ*1, *ΔphaZ*2, and *ΔphaZ1 ΔphaZ2* using ordinary one-way ANOVA followed by Tukey-adjusted pairwise comparisons, with each deletion strain compared with WT.

For the complementation experiment, PHA content was compared across the seven strains by Brown-Forsythe and Welch ANOVA with Dunnett’s T3 multiple-comparisons test, as it does not assume equal variances. Gompertz-derived growth parameters were compared by ordinary one-way ANOVA with Šídák’s correction; homogeneity of variance was assessed by the Brown-Forsythe test and was not rejected.

The effects of genotype and nutrient limitation on PHA accumulation were evaluated using ordinary two-way ANOVA. Genotype and nitrogen or phosphorus availability were treated as fixed factors, and the full model included the genotype-by-nutrient interaction. When a significant interaction was detected, WT and *ΔphaZ1 ΔphaZ2* were compared within each nutrient condition using Šídák’s multiple-comparisons test. Nitrogen-free and severely phosphorus-limited conditions that did not produce sufficient biomass for PHA quantification were excluded from the corresponding PHA analyses.

The effects of genotype and cultivation strategy on PHA content were also evaluated using ordinary two-way ANOVA, with genotype and cultivation strategy as fixed factors and with inclusion of the genotype-by-cultivation interaction. Šídák’s multiple-comparisons test was used for comparisons between genotypes within each cultivation strategy and between one-stage and two-stage cultivation within each genotype. All reported P values from post hoc comparisons were adjusted for multiple testing. Potential outliers were evaluated by the Grubbs test (α= 0.05) and the ROUT method (Q = 1%); no data points were excluded from any analysis.

## Data availability

Whole-genome sequencing reads for the ΔphaZ2 and ΔphaZ1 ΔphaZ2 strains have been deposited in the NCBI Sequence Read Archive under BioProject accession PRJNA1541609. Raw growth, LC–MS, and FTIR data underlying all figures, the LC–MS calibration curves and limit-of-detection calculations, the FTIR calibration set and per-sample predictions, and the derivation of the OD₆₆₀-to-dry-cell-weight conversion factor are provided in the supplemental material. Analysis scripts for modified Gompertz growth-curve fitting, FTIR–LC–MS calibration, and PHA prediction from FTIR spectra are archived at Zenodo (https://doi.org/10.5281/zenodo.23274022).

## Acknowledgments

We would like to thank Brian Gallagher III and Hailee Morrisson for guidance with culture techniques and contamination troubleshooting. We also thank Gavin Yao for early work toward the Rpal_0568 deletion, which informed the cloning strategy used in this study.

The following grants to AB supported this work: National Science Foundation (NSF) grants 2021822, 2124088, 2117198, and 2300081; Defense Established Program to Stimulate Competitive Research (DEPSCoR) grant FA9550-21-1-0211; and a David & Lucile Packard Foundation Fellowship. JB is supported by National Institute of General Medical Sciences (NIGMS) grant NIH 1R35GM158225.

