## Supplemental Materials for "Deletion of two candidate depolymerases with overlapping genetic effects increases PHA yield in *Rhodopseudomonas palustris* TIE-1"

Table S1. Primers used in the study

| **Primer** | **Sequence (5′→3′)** | **Purpose** |
| --- | --- | --- |
| **Deletion construct (*Rpal_0568, phaZ2*)** |  |  |
| TC1 | taattccctagtggatcccccACCAGCCAGAAC  GAGTTGAACAGC | Upstream homology arm, forward |
| TC7 | AAGTTGCGACCGTCCTGGTCTGGGCCCCTGC | Upstream homology arm, reverse |
| TC8 | GACCAGGACGGTCGCAACTTCCGACTGCC | Downstream homology arm, forward |
| TC4 | ttgatatcgaattcctgcagcccTTCCTGGCCAAT  CAGCTGAATTGCAGC | Downstream homology arm, reverse |
| TC5 | gggctgcaggaattcga | pJQ200KS vector, forward |
| TC6 | gggggatccactaggga | pJQ200KS vector, reverse |
| **Genotyping and screening (*Rpal_0568, phaZ2*)** |  |  |
| TC9 | CAGGATGCCGAACCACATCAGG | Right upstream of deletion |
| TC10 | GCCAACAACGAAGCTCCGCAG | Right downstream of deletion |
| TC11 | GCAAAGACCAGCACGCGCCA | screening, upstream |
| TC12 | TCGCCGTCACGTGCACCAC | screening, downstream |
| TC13 | ATCTTCGCCACCGGCCTGTC | Internal forward |
| TC14 | GTGCTACGTCCGTGGTCAGTTC | Internal reverse |
| **Genotyping and screening (*Rpal_0578, phaZ1*)** |  |  |
| TC15 | CCCAACTGCTTCAACCTG | screening, upstream |
| TC16 | ATAGCCGCAGACCTCGTTCT | screening, downstream |
| TC17 | CGAATTCGTCCACGATGAG | Internal forward |
| TC18 | TCGTCGACACCGAGAATG | Internal reverse |
| **Complementation constructs** |  |  |
| TC22 | Tttcacacaggaaacagcata  **ATGCCGGTTGTCATTGGTGAATTT** | *phaZ1*, forward (single and tandem) |
| TC23 | cttggcgtaatcatggtca**TCACTCCGCCGCGC** | *phaZ1*, reverse (single) |
| TC24 | tttcacacaggaaacagcata**GTGTCACT**  **TAAGAATCCTATCGACTTCCTCG** | *phaZ2*, forward (single) |
| TC25 | cttggcgtaatcatggtca**CTACTTCATCAATC**  **CGGCAGCGG** | *phaZ2*, reverse (single and tandem) |
| TC26 | GA*actgtagAGGAAAtatcaga***GTGTCACTT**  **AAGAATCCTATCGACTTCCTCG** | *phaZ2*, forward (tandem; carries linker) |
| TC27 | AC*tctgataTTTCCTctacagt***TCACTCCGCCGCGC** | *phaZ1*, reverse (tandem; carries linker) |

Lowercase letters indicate 5′ overhangs homologous to the vector or, in TC26 and TC27, the linker and ribosome binding site.

Figure S1.
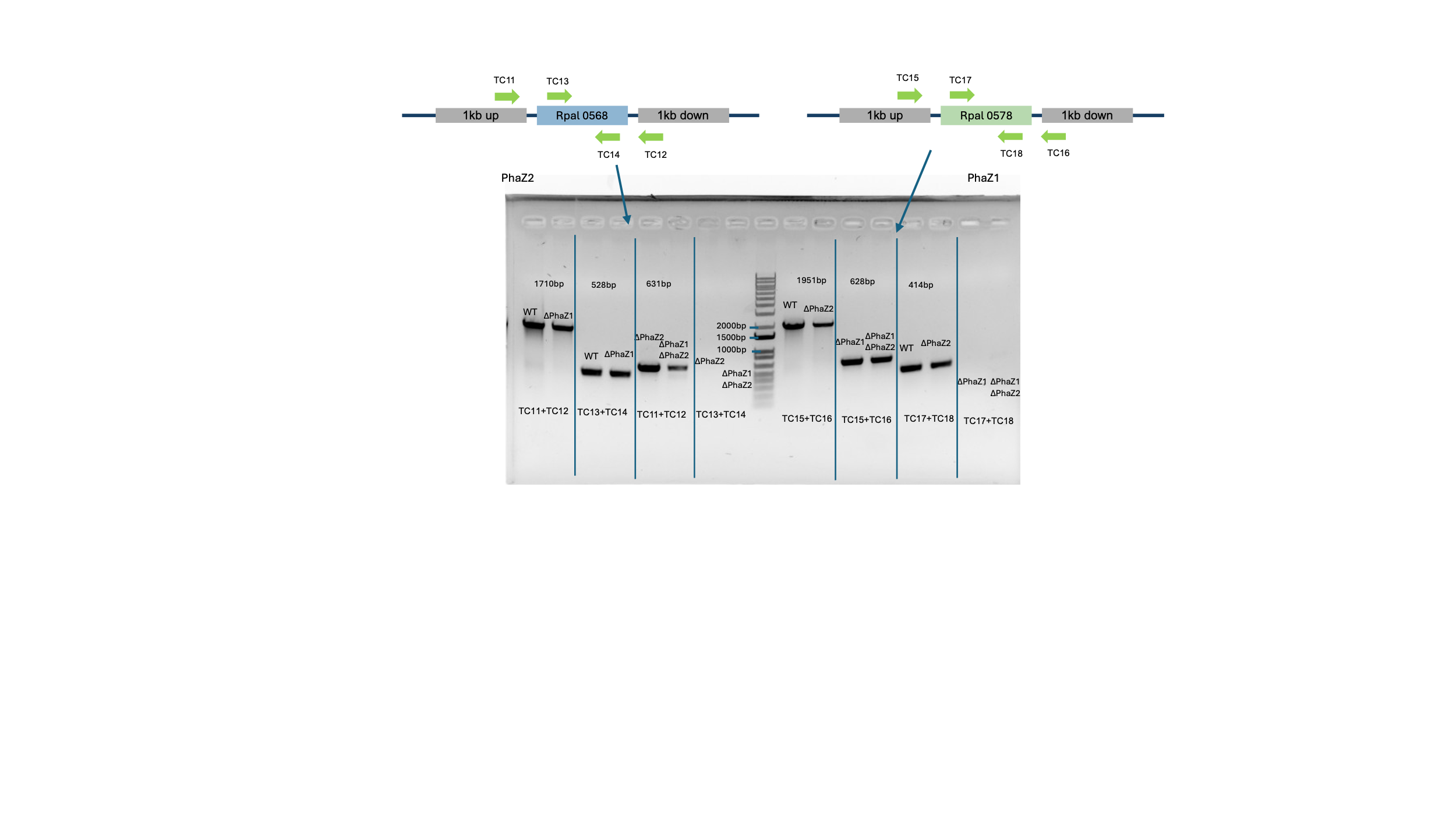

**Figure S1 | PCR verification of markerless Rpal_0568 and Rpal_0578 deletions.** (Top) Primer positions at the Rpal_0568 (*phaZ2*) and Rpal_0578 (*phaZ1*) loci. Flanking primers (TC11/TC12, TC15/TC16) anneal outside the approximately 1-kb homology arms; internal primers (TC13/TC14, TC17/TC18) anneal within each coding sequence. Primer sequences are given in Table S1. (Bottom) Genomic PCR on wild-type TIE-1 and the Δ*phaZ1*, Δ*phaZ2*, and Δ*phaZ1* Δ*phaZ2* strains. Flanking primers yielded 1,710 bp at *phaZ2* and 1,951 bp at *phaZ1* in strains retaining the locus, and 631 bp and 628 bp respectively in strains carrying the deletion, consistent with removal of 1,079 bp of Rpal_0568 and 1,323 bp of Rpal_0578. Internal primers yielded 528 bp (*phaZ2*) and 414 bp (*phaZ1*) only in strains retaining the corresponding gene, and no product in the deletion strains. Because the same genomic DNA preparations gave the expected flanking-primer amplicons, the absence of an internal product reflects loss of the coding sequence rather than failed amplification. All lanes are from a single gel; vertical lines mark groupings by primer pair. Deletions were additionally confirmed by whole-genome sequencing.

Figure S2.

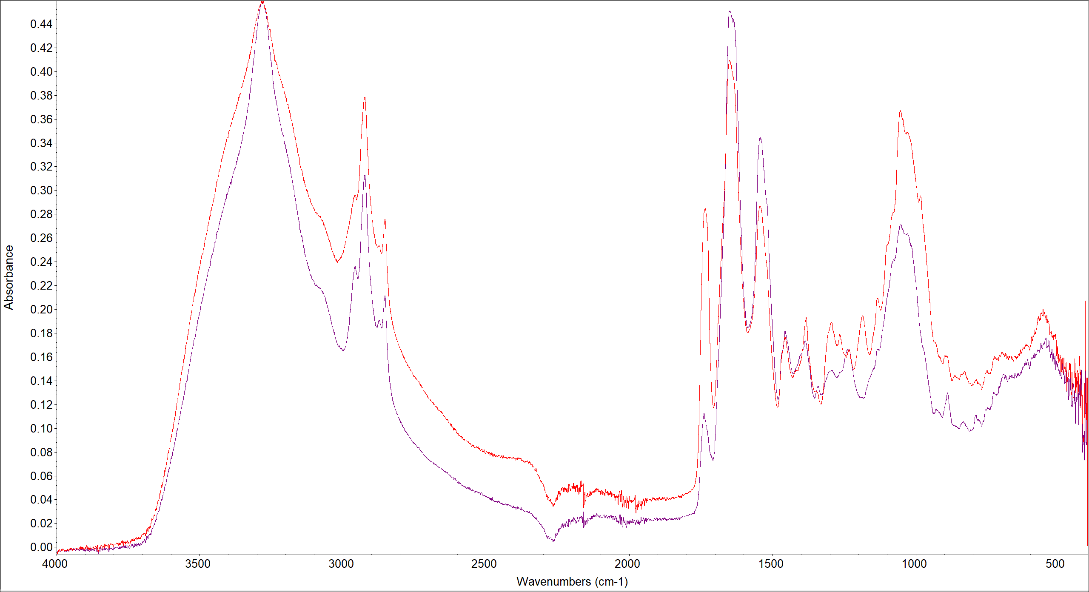

25% nitrogen

*ΔphaZ1 ΔphaZ2*:

strong PHA accumulation

100% nitrogen WT:

weak PHA accumulation

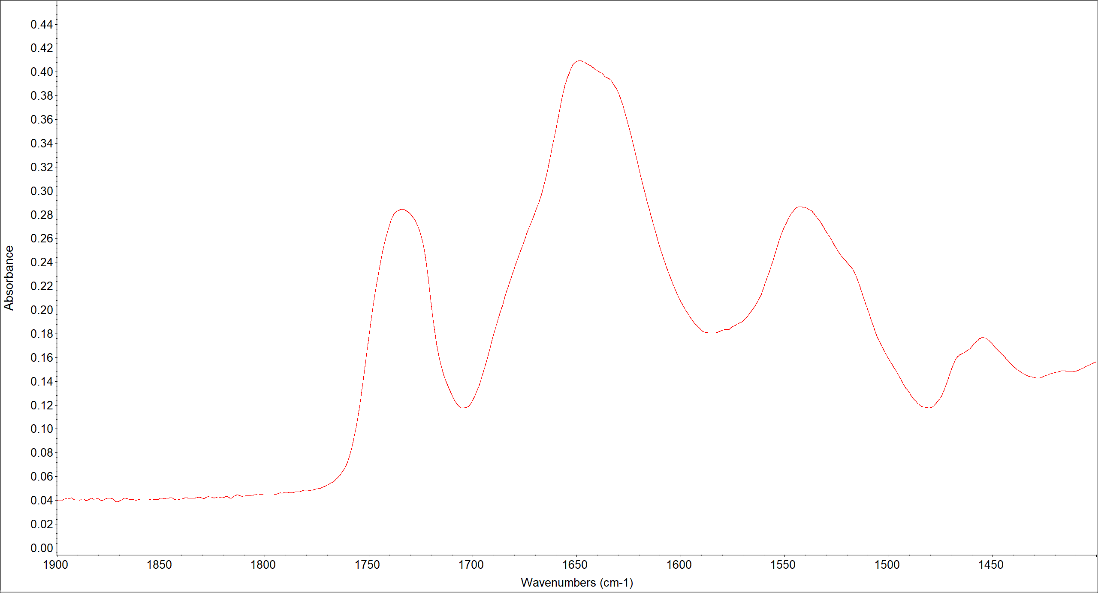

PHA Carbonyl:

1700-1750 cm**^-1^**

Biomass Amide I:

1600-1700 cm**^-1^**

Biomass Amide II:

1500-1580 cm**^-1^**

**A**

**B**

**Figure S2 | LC–MS-calibrated FTIR analysis of TIE-1.** (A) Representative FTIR spectra of a high-PHA sample (*ΔphaZ1 ΔphaZ2* grown with 25% nitrogen) and a low-PHA sample (WT grown with 100% nitrogen). The shaded region marks the PHA-associated carbonyl band. (B) Spectral regions integrated for the PHA carbonyl signal (1700–1750 cm⁻¹), amide I signal (1600–1700 cm⁻¹), and amide II signal (1500–1580 cm⁻¹). The carbonyl peak area was normalized to the combined amide I and amide II areas.

Figure S3.

**A**

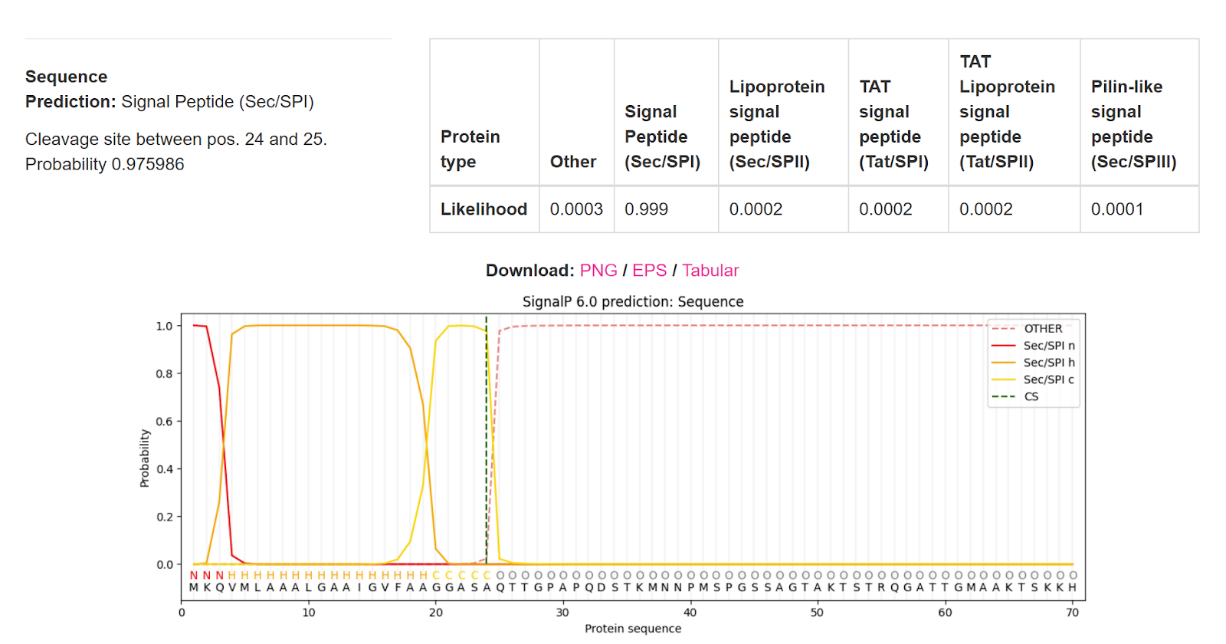

**B**

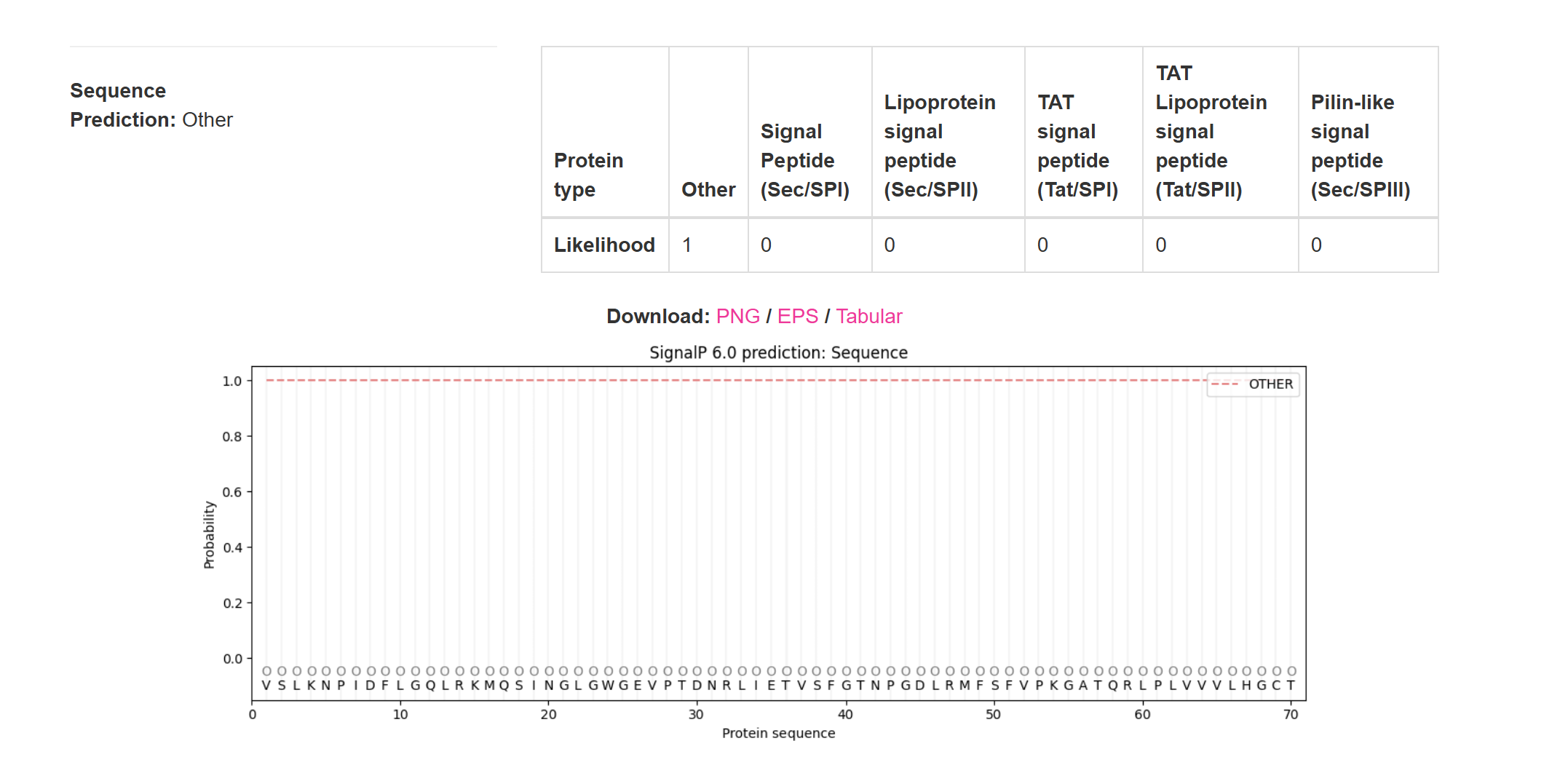

**C**

Signal peptide upstream of PhaZ2

PhaZ2

- Stop codon

MKQVMLAAALGAAIGVFAAGGASAQTTGPAPQDSTKMNNPMSPGSSAGTAKTSTRQGATTGMAAKTSKKHPTKKNDVQPSSMSKGSAK-LAFAEFVHDDGPPPPSCGPFCGWLRADPITVQGLPYCNAAIIWKSGQGPRPGAT

VSLKNPIDFLGQLRKMQSINGLGWGEVPTDNRLIETVSFGTNPGDLRMFSFVPKGATQRLPLVVVLHGCTQNAAAYEIGAGWCTLAQRYGFALLLPEQTRTNNPNTCFNWFSPDDIRRGSGEVGSIHQMIEHMTRTHQIDQARIFATGLSAGGGMTAALLATYPDVFAGGAVIAGLPYGVAHNVRQALKEMRHATPRSGPELGELVRHATAHQGPWPKLAVWHGDADHTVDPANADALVRQWLDLHGLAEAPTAQNEVCGYPHRAWRNSSGETVVESYTVTGMAHGTPLGAGAGDERYGVAGAYLLDVGISSSYHIAQFFGLTSRIHQPRSADTSAKDRPAHDGHHELTTDVARRSRKSAGTGHAEGAAAPPHGKRGIDVGAVITRALTAAGLMK-

**Fig. S3** **The Rpal_0568 locus of *Rhodopseudomonas palustris* TIE-1 encodes a signal peptide that is separated from the depolymerase coding sequence.** **(A)** SignalP 6.0 analysis of the 88-codon open reading frame immediately upstream of *phaZ2*. The sequence is assigned to the Sec/SPI class with a likelihood of 0.999, with cleavage predicted between positions 24 and 25. Colored traces give the per-residue probability assigned to each region of a Sec/SPI signal peptide — the positively charged n-region, the hydrophobic h-region, and the c-region containing the Ala-X-Ala cleavage motif — and to the mature-protein class (OTHER); the vertical dashed line marks the predicted cleavage site. **(B)** SignalP 6.0 analysis of the 395-residue PhaZ2 product, which is assigned to the OTHER class with a likelihood of 1.00; no signal peptide of any class is predicted. **(C)** Conceptual translation of the locus. Red, the 88-codon upstream open reading frame, of which the first 24 residues constitute the predicted signal peptide; black, the 54 codons separating the two reading frames; blue, the 395-residue PhaZ2 coding sequence. Dashes mark stop codons. The upstream open reading frame terminates at an in-frame stop codon, so the signal peptide and the depolymerase coding sequence cannot be translated as a single polypeptide. The stop codon was confirmed by whole-genome sequencing of TIE-1 in this study and is not an artifact of the reference assembly. Predictions were made with SignalP 6.0 using default parameters.

**Table S2.** Protein sequences used in the phylogenetic analysis of TIE-1 PHA depolymerase candidates.

| **Accession (residues)** | **Organism** |  | **Length in alignment** | **Clade (Fig. 1A)** |
| --- | --- | --- | --- | --- |
| Rpal_0568 | *Rhodopseudomonas palustris TIE-1 (PhaZ2)* |  | 381 | Extracellular type 1 |
| WP_011156133.1 (16–395) | *Rhodopseudomonas palustris CGA009* |  | 380 | Extracellular type 1 |
| WP_107342770.1 (16–394) | *Rhodopseudomonas palustris* |  | 379 | Extracellular type 1 |
| WP_047307393.1 (16–385) | *Rhodopseudomonas palustris* |  | 370 | Extracellular type 1 |
| WP_114356866.1 (16–398) | *Rhodopseudomonas sp. (MULTISPECIES)* |  | 383 | Extracellular type 1 |
| WP_407167513.1 (16–393) | *Bradyrhizobium sp. ORS 111* |  | 378 | Extracellular type 1 |
| WP_426438009.1 (16–393) | *Bradyrhizobium genosp. P* |  | 378 | Extracellular type 1 |
| WP_044586480.1 (16–393) | *Bradyrhizobium sp. LTSPM299* |  | 378 | Extracellular type 1 |
| Rpal_0578 | *Rhodopseudomonas palustris TIE-1 (PhaZ1)* |  | 442 | Intracellular |
| WP_319942045.1 | *Rhodopseudomonas palustris* |  | 442 | Intracellular |
| WP_110782234.1 | *Rhodopseudomonas faecalis* |  | 442 | Intracellular |
| WP_184257698.1 | *Rhodopseudomonas rhenobacensis* |  | 442 | Intracellular |
| WP_184084816.1 | *Afipia massiliensis* |  | 441 | Outgroup |
